# Optimized nickase- and nuclease-based prime editing in human and mouse cells

**DOI:** 10.1101/2021.07.01.450810

**Authors:** Fatwa Adikusuma, Caleb Lushington, Jayshen Arudkumar, Gelshan I. Godahewa, Yu C.J. Chey, Luke Gierus, Sandra Piltz, Daniel Reti, Laurence OW. Wilson, Denis C. Bauer, Paul Q. Thomas

## Abstract

Precise genomic modification using prime editing (PE) holds enormous potential for research and clinical applications. Currently, the delivery of PE components to mammalian cell lines requires multiple plasmid vectors. To overcome this limitation, we generated all-in-one prime editing (PEA1) constructs that carry all the components required for PE, along with a selection marker. We tested these constructs (with selection) in HEK293T, K562, HeLa and mouse embryonic stem (ES) cells. We discovered that PE efficiency in HEK293T cells was much higher than previously observed, reaching up to 95% (mean 67%). The efficiency in K562 and HeLa cells, however, remained low. To improve PE efficiency in K562 and HeLa, we generated a nuclease prime editor and tested this system in these cell lines as well as mouse ES cells. PE-nuclease greatly increased prime editing initiation, however, installation of the intended edits was often accompanied by extra insertions derived from the repair template. Finally, we show that zygotic injection of the nuclease prime editor can generate correct modifications in mouse fetuses with up to 100% efficiency. In summary, PE-nuclease and the PEA1 plasmids provide new tools to generate intended edits with high efficiency.

## INTRODUCTION

CRISPR prime editing (PE) is a versatile genome editing technology that enables the introduction of precise sequence modifications including specific indels and all types of point mutations. The prime editor complex consists of an SpCas9 nickase (H840A)-Reverse Transcriptase (RT) fusion protein and a prime editing gRNA donor template (pegRNA) comprising a gRNA sequence with a 3’ extension encoding the desired edit [1]. Binding and reverse transcription of the pegRNA enables the desired edit to be introduced into the nicked strand at the target site. Subsequent flap equilibration, cleavage and mismatch repair leads to permanent incorporation of the edit. The addition of another gRNA to create a second nick on the opposite strand significantly improves PE efficiency, a strategy termed PE3 [1].

PE has been shown to create specific edits with unprecedented and relatively high efficiency in HEK293T cells [1]. However, PE was reported to be less efficient in other cell types such as HeLa, K562 and U2OS [1]. Importantly, the multi-vector approach used by Anzalone et al. to deliver the PE components did not include selection of transfected cells [1]. Since transfection is required for editing, PE efficiency may have been underestimated in this study.

To address this issue, we generated Prime Editing All-in-One (PEA1) plasmids consisting of three cassettes for expression of all PE3 components and a selection marker. We show that PEA1 editing constructs can be readily generated using a one-step golden gate digestion-ligation protocol [2]. Using PEA1 constructs with selection, we performed PE in HEK293T, K562, HeLa and mouse ES cells, which revealed that the efficiency of PE is generally higher than previously estimated. We also showed that Cas9 nuclease (as opposed to Cas9 nickase) could further improve PE efficiency in cultured cells and in mouse zygotes, where up to 100% of mouse fetuses were correctly edited.

## RESULTS

### Efficient one-step generation of prime editing constructs using an all-in-one plasmid system

We previously developed an all-in-one plasmid system that enables rapid generation of dual gRNA expression constructs [2]. Using the same principle, we generated all-in-one PE constructs to simplify PE experiments. Our Prime Editing All-in-One (PEA1) plasmids contain all of the components for PE3 editing, as well as a puromycin or GFP selection marker coupled to the prime editor using a T2A self-cleaving peptide (PEA1-Puro and PEA1-GFP, respectively) (Fig. 1a). PEA1 also contains three BbsI golden gate cloning sites (Fig. 1a, and Supplementary Fig. 1) to enable the simultaneous insertion of annealed oligonucleotides encoding the gRNA protospacer, the RT template and the second nick gRNA protospacer using a one-step digestion-ligation protocol [2]. Additionally, we are developing an online bioinformatics tool called PETAL (Prime Editing Target Locator) to facilitate PE experiments. A beta version of PETAL is available online which we are continuing to develop. PETAL is intended to help users select the gRNA, RT template and second-nick gRNA sequences, and design the oligonucleotides required for the generation of the corresponding constructs, including our PEA1 system (Supplementary Fig. 2). We tested the efficiency of PEA1 construct generation by generating 24 PE constructs equivalent to those used by Anzalone et al. to edit *HEK3, RNF2, RUNX1*, and *VEGFA*, using identical pegRNA and second-nick gRNA sequences. From 66 plasmids screened by restriction enzyme digestion (Supplementary Fig. 3), 48 constructs (72.7%) contained complete oligonucleotide duplex integration. Thus, PEA1 plasmids, combined with the one-step digestion-ligation protocol, enable efficient generation of all-in-one PE constructs of interest.

**Figure 1.**
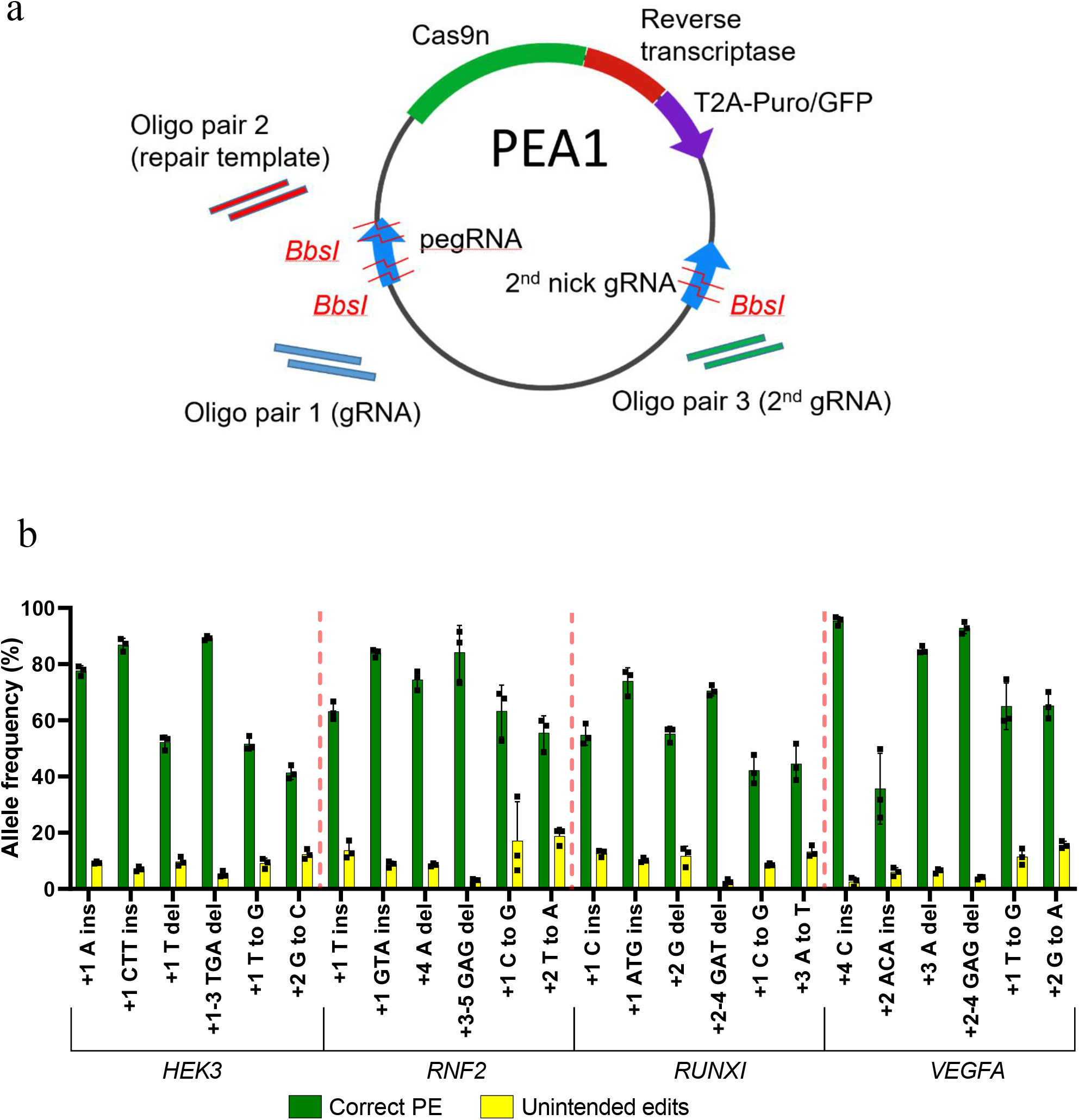
Prime editing using an all-in-one plasmid vector in HEK293T cells. (a) Schematic representation of the PEA1 plasmid, containing three BbsI golden gate assembly sites to insert the customizable guide target sequences and RT template. (b) PEA1 editing efficiencies of targeted 1- and 3-bp insertions, 1- and 3-bp deletions and point mutations at four genomic sites in HEK293T cells. Mean ± SD, n=3.

### Enhanced Prime editing outcomes in HEK293T cells

To assess the editing efficiency of the PE3 approach using the PEA1-Puro system, we transfected each of the 24 constructs into HEK293T cells and selected for transfected cells using puromycin. Deep amplicon NGS showed that PE efficiency was very high with average correct PE efficiency of 67% (Fig. 1b), exceeding that observed by Anzalone et al. for all target sites [1]. Twenty PEA1-Puro constructs generated correct prime edits with >50% efficiency, and 11 of them exceeded 70% efficiency (Fig. 1b). The highest editing efficiency was 95% (VEGFA +4 C Ins) (Fig. 1b). In contrast, Anzalone et al., who performed PE3 experiments using multiple vectors and without selection, showed ~35% average PE efficiency across the 24 edits with only 2 correct prime edits with >50% efficiency and none with >70% efficiency (Fig 4a, c, d & f of Anzalone et al.) [1]. These data indicate that PE3 efficiency shown by Anzalone et al. in HEK293T cells, although relatively high, was likely an underestimation of the actual PE activity, which can be further enhanced using our PEA1 plasmid system and a selection step.

### Prime editing is less efficient in K562 and HeLa cells

Despite the high efficiency of PE in HEK293T cells, Anzalone et al. observed that PE3 was generally less efficient in other cell lines such as K562 and HeLa [1]. It is unclear whether this was due to lower transfection efficiency or a reduced propensity for PE repair. To investigate this, we used our PEA1-Puro constructs and puromycin selection protocol to assess the efficiency of the PE3 approach in K562 and HeLa cells. Twelve of the previously generated PEA1-Puro constructs were selected for analysis, including one insertion, one deletion and one point mutation for each of the four target sites. Four of these edits (HEK3 CTT ins, HEK3 T to G, RNF2 GTA ins, and RNF2 C to G) were previously tested by Anzalone et al. in K562 and HeLa cells with multiple vectors without selection [1]. In K562, we showed 26%, 33%, 48% and 44% correct PE efficiency, respectively, for these edits (Fig. 2a). In comparison, Anzalone et al. observed 25%, 21%, 7.8% and 5% correct PE efficiency, respectively (Fig. 5e and Extended Fig. 10c of Anzalone et al.) [1]. In HeLa cells, our correct PE efficiency for these edits was 5.3%, 9.7%, 17.7% and 11.7%, respectively (Fig. 2b), compared to 12%, 12.6%, 4.6% and 3.6%, respectively (Fig. 5e and Extended Fig. 10e of Anzalone et al.) [1]. This indicates that PE3 using PEA1 and a selection step could improve the PE efficiency in these cell lines. However, this improvement did not match the PE efficiencies observed in HEK293T cells (Fig. 1b). The average correct PE efficiency for these 12 edits was 71% in HEK293T cells, yet only 29% in K562 and 7.2% in HeLa cells. Together, this indicates that while PE3 is highly efficient in HEK293T cells, its efficiency is moderate in K562 cells and low in HeLa cells, which may reflect differences in DNA repair activity in these cell lines.

**Figure 2.**
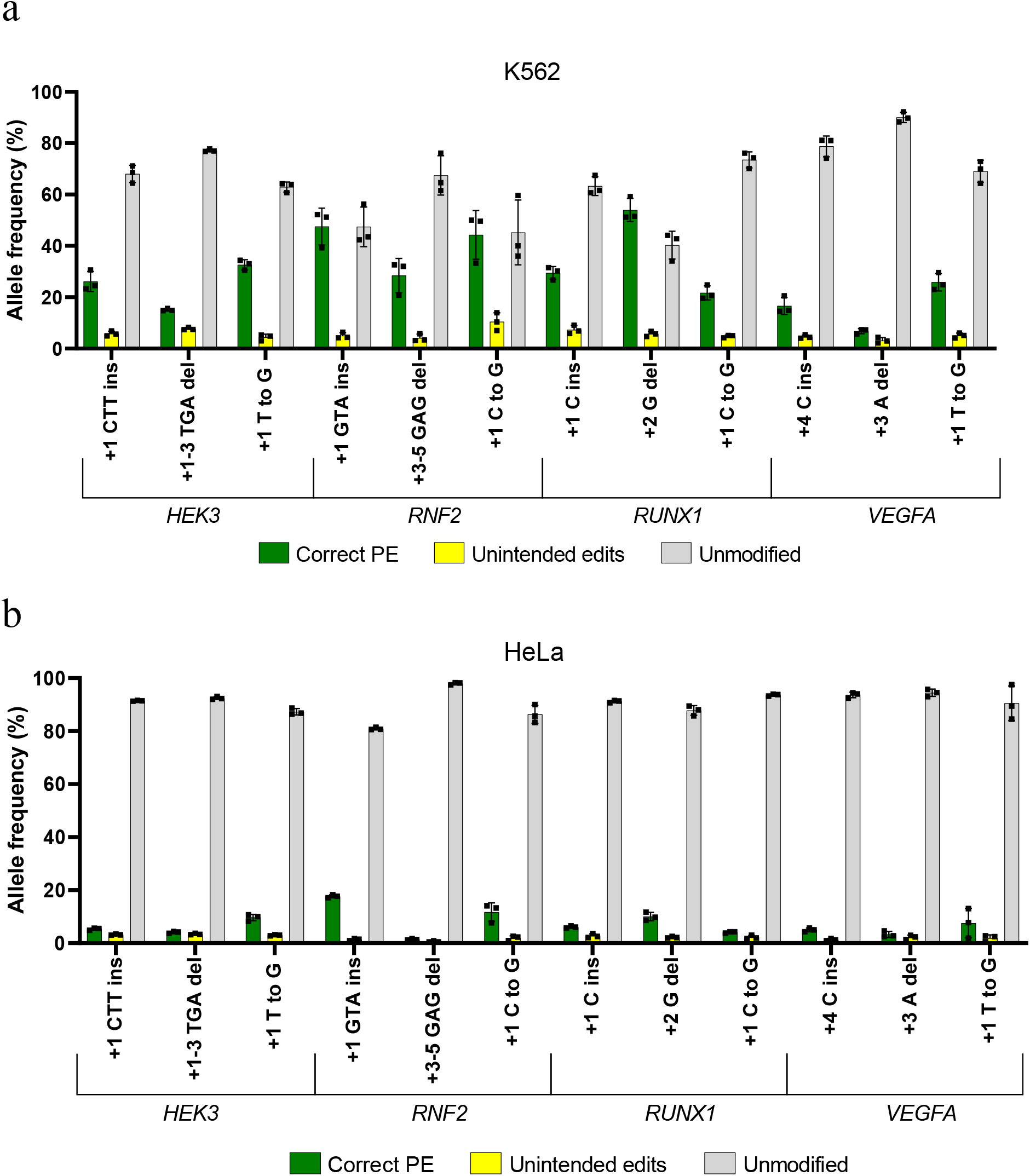
Prime editing using an all-in-one plasmid vector in hard-to-edit cell lines. PEA1 editing efficiencies of targeted 1- or 3-bp insertions, 1- or 3-bp deletions and point mutations in (a) K562 and (b) HeLa cell lines. Mean ± SD, n=3.

### Prime editing using nuclease prime editor in HeLa and K562 cells

We hypothesized that the low PE efficiencies observed in HeLa and K562 cells might be affected by inefficient 5’ flap resection and removal of the non-edited DNA strand, which conceivably restricts its replacement. We reasoned that using SpCas9-nuclease instead of the SpCas9 nickase (H840A) bypasses this requirement. Thereby, we generated PE nuclease constructs (PEA1-Nuc) by replacing the H840A nickase sequence in PEA1 with the WT SpCas9 nuclease sequence. We then generated 12 PEA1-Nuc-Puro editing constructs equivalent to the PEA1-Puro constructs that we tested in K562 and HeLa cells. Since the SpCas9 nuclease creates a double-stranded break, the second-nick gRNA required for the PE3 system was not used. To assess editing efficiency in K562 and HeLa cells, the 12 PEA1-Nuc-Puro constructs were transfected, followed by puromycin selection. Surprisingly, although correct PE was still low, averaging 22% and 6.7% in K562 and HeLa cells, respectively, the NGS analysis revealed that the frequency of alleles incorporating any intended edits was very high, averaging 71% and 30% in K562 and HeLa, respectively (Fig. 3a & b). Unexpectedly, the majority of alleles containing any intended edits were found to have extra sequences derived from the RT template (Fig. 3c). These extra sequences likely resulted from insertion of RT template sequences at the unmodified break sites, thus installing the intended edits with partial duplication of the template sequences (Fig. 3c). These partial template duplications (PTDs) were presumably formed due to imperfect resolution of PE events in which the reverse transcribed templates underwent end-joining with the downstream DNA junction at the pegRNA break sites instead of annealing with the genomic homology sequences (Supplementary Fig. 5). PTD frequencies averaged 49% and 23% in the PE-nuclease-treated K562 and HeLa cells, respectively (Fig. 3a & b). This indicates that the nuclease prime editor could enhance the prime editing initiation (priming and reverse transcription) in K562 and HeLa cells. However, in most cases, correct resolution was not achieved and generated PTDs instead of the correct PE. Unsurprisingly, the nuclease prime editor also induced a higher level of indels and greatly reduced the proportion of the unmodified alleles (Fig. 3a & b).

**Figure 3.**
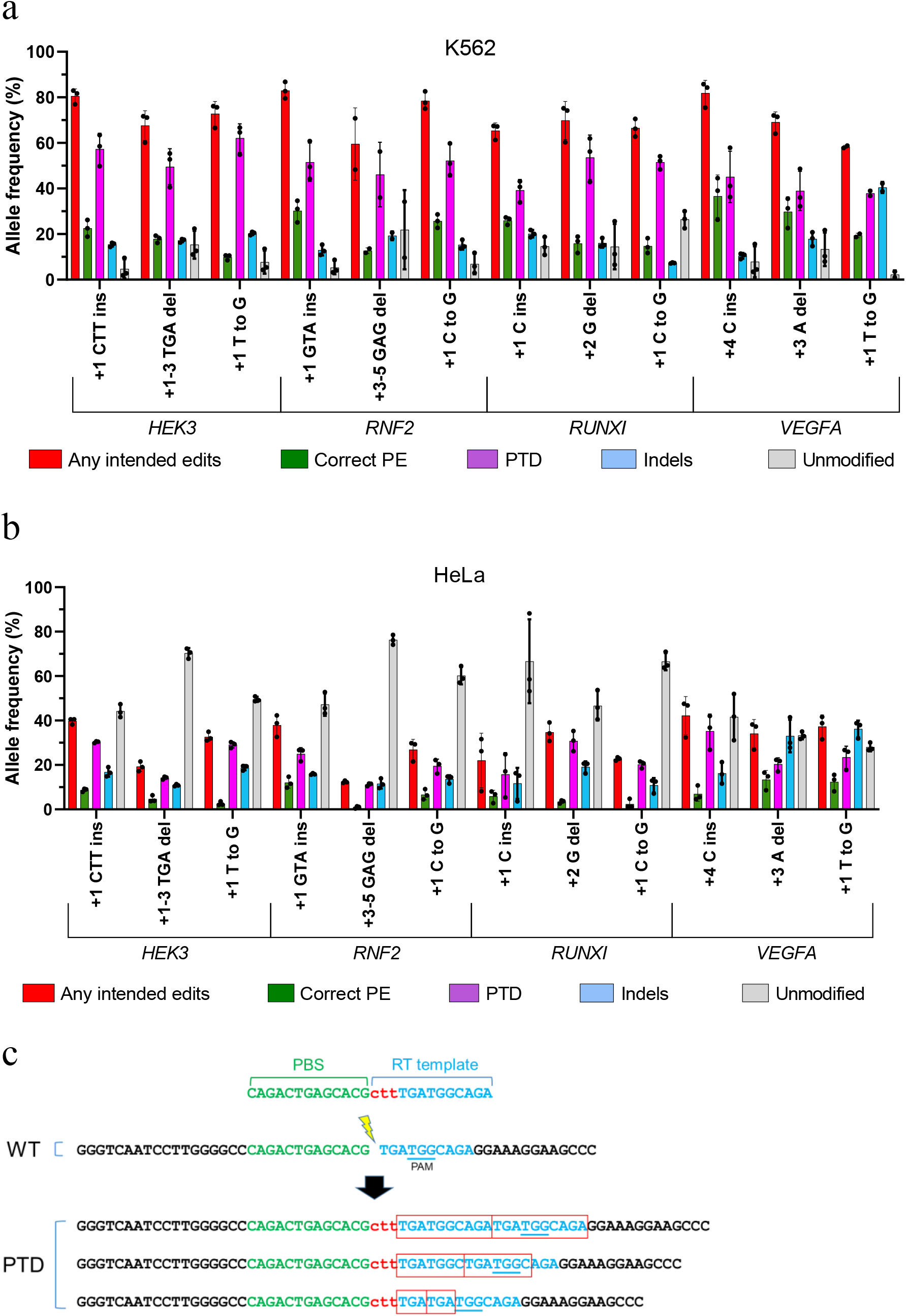
Nuclease prime editing in hard-to-edit cell lines. PEA1-Nuc editing efficiencies of targeted 1- or 3-bp insertions, 1- or 3-bp deletions and point mutations in (a) K562 and (b) HeLa cell lines. Mean ± SD, n=3, except for RNF2 +3-5 GAG del and VEGFA +1 T to G in K562 which were n=2. (c) Example of partial template duplications (PTDs) observed in HEK3 +1 CTT ins nuclease prime edited cells.

We also characterized whether PTDs were also generated by the PE3 nickase. Unexpectedly, we found that PTDs were present at all 24 PE target sites tested in HEK293T cells and comprised 5-40% of the unintended edits (Supplementary Fig. 4).

### Prime editing in mouse ES cells using PEA1 and PEA1-nuclease

We also tested PE3 and PE-nuclease approaches in mouse ES cells using PEA1-puro and PEA1-Nuc-Puro, respectively. Nine different small edits including 3 nt insertions, point mutations, 3 nt deletions were tested for each system across 4 loci. Although the efficiency was variable, correct PE editing was induced at all sites using PE3 or PE-nuclease (Fig. 4a & b). We observed considerable variation in the editing efficiency. For example, PE3 resulted in 85% and 74% correct editing when creating +1 CTC ins and +5 G to C edits at the *Chd2* target site, respectively (Fig. 4a). In contrast, PE3 editing efficiency was 19% and 1.7% when creating +1 TGT ins and +6 G to A edits at the *Tyr* target site, respectively (Fig. 4a). PE3 at the *Mixl1* target site resulted in very low correct PE efficiency when using second-nick gRNA +48, but the efficiency was higher when using second-nick gRNA - 60 (Fig. 4a), indicating that optimizing the position of the second-nick gRNA can contribute to higher PE efficiency using the PE3 approach.

**Figure 4.**
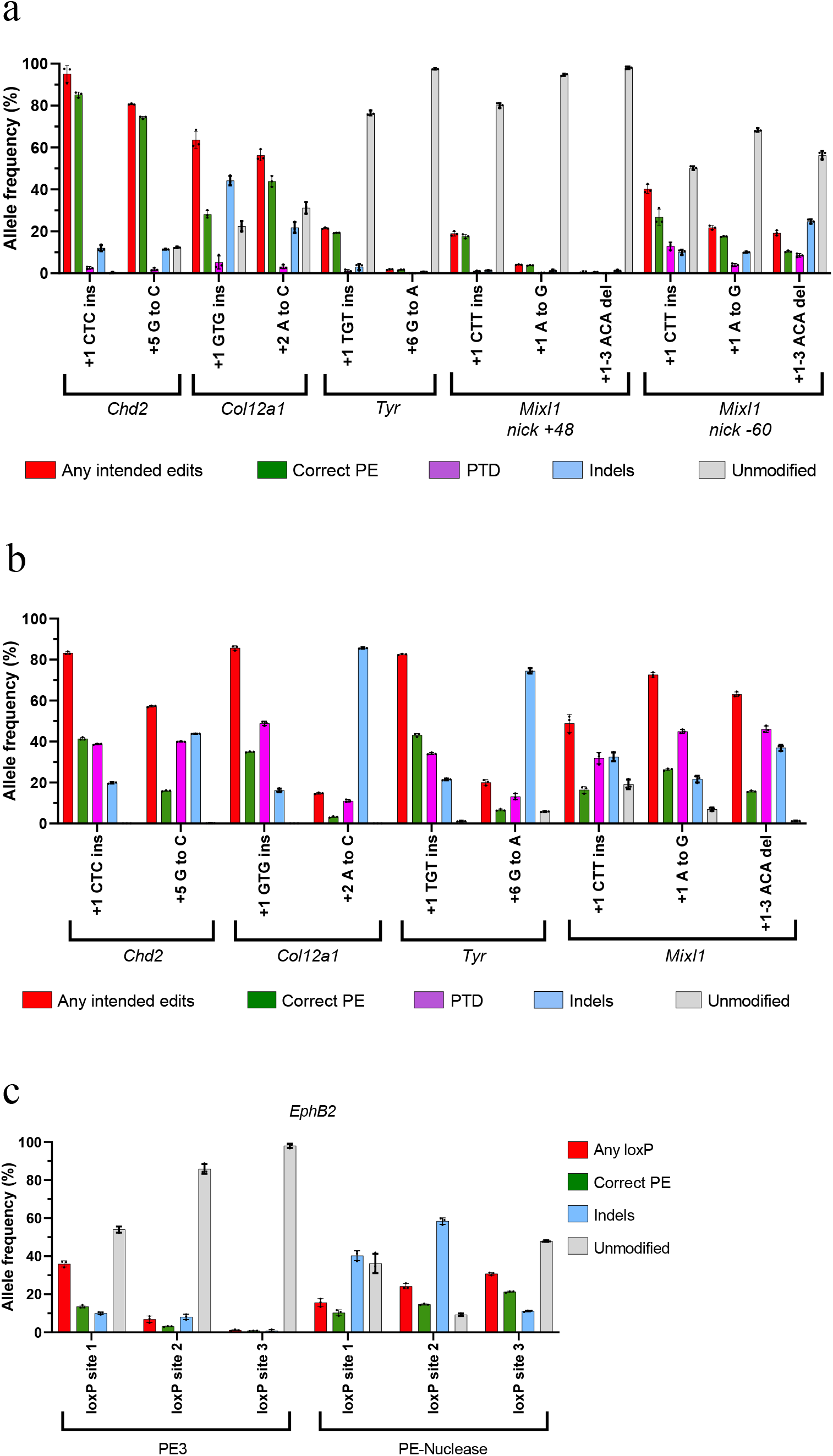
Prime editing PE3 and PE-nuclease in mouse embryonic stem (ES) cells. Creating small edits using PEA1 (a) or PEA1-Nuc (b) vector. (c) Inserting loxP sequences into the *EphB2* locus using PEA1 and PEA1-Nuc Mean ± SD, n=3. ‘Any loxP’ was defined as alleles containing the full length of loxP sequences with or without extra modifications.

Similar to our previous findings, PE-nuclease produced a high frequency of alleles with the installation of the intended edits predominated by PTDs. It also induced a higher level of indels and lower unmodified alleles (Fig. 4b). Interestingly, for 5 of the 9 edits, PE-nuclease induced higher correct PE efficiency than the PE3. For example, at the *Tyr* target site, PE-nuclease induced 43% and 6.6% correct PE efficiency when creating +1 TGT ins and +6 G to A edits, respectively, which were 2-3 times higher than the efficiency induced by PE3 (Fig. 4a & b). However, there were some target site at which the PE-nuclease generated much lower correct PE efficiencies compared to PE3, such as *Chd2* +1 CTC ins (41% vs 85% for PE-nuclease vs PE3, respectively) and *Chd2* +5 G to C (16% vs 74% for PE-nuclease vs PE3, respectively; Fig. 4a & b). Although the PE-nuclease induced higher correct editing for *Col12a1* + 1 GTG ins (35% vs 28% for PE-nuclease vs PE3, respectively), PE-nuclease efficiency dropped to 3.2% when creating +2 A to C edit, which was much lower than PE3 (44%; Fig. 4a & b). We suspect that the low efficiency of the +2 A to C edit using PE-nuclease was due to re-cutting, as the 1-nt substitution was located in the gRNA targeting sequence [3, 4]. Consistent with this, we noticed a high frequency of prime edited alleles with indels (Supplementary Fig. 6), which were rare for PE3 *Col12a1* + 2 A to C. This suggests that PE-nuclease is more prone to re-cutting, particularly when the editing is relatively subtle (e.g. point mutations). We also attempted to insert longer loxP sequences (40 nt) at 3 different intronic sites in *EphB2* locus. While PE3 induced relatively efficient loxP insertions in 1 of 3 sites tested, PE-nuclease consistently induced efficient insertions at all 3 sites (Fig. 4c).

### Efficient generation of mice with specific edits by zygotic injection of nuclease prime editor

The PE3 system has previously been used to generate mice with specific edits by zygote microinjection [5, 6]. However, the efficiency was low, and installation of the PE mutations often induced large deletions due to concurrent dual nicking. Although PE2 reduced indel generation, on-target editing efficiency was also decreased [5, 7]. Given the relatively high efficiency of PE-nuclease in cell culture experiments, we sought to assess PE-nuclease efficiency in microinjected zygotes. We initially tested this by generating 3 nt insertions at the *Chd2* and *Col12a1* sites that were previously targeted by Aida et al. using identical PBS and homology arms [5]. Remarkably, these 3 nt insertions were generated with very high efficiency; 21 of 24 mice (87.5%) and 6 of 7 mice (85.7%) carried the correct insertions at *Chd2* and *Col12a1*, respectively (Fig. 5). We then attempted to generate a 3 nt insertion at the *Tyr* site and observed the correct edit in all 19 mice (100%) (Fig. 5).

**Figure 5.**
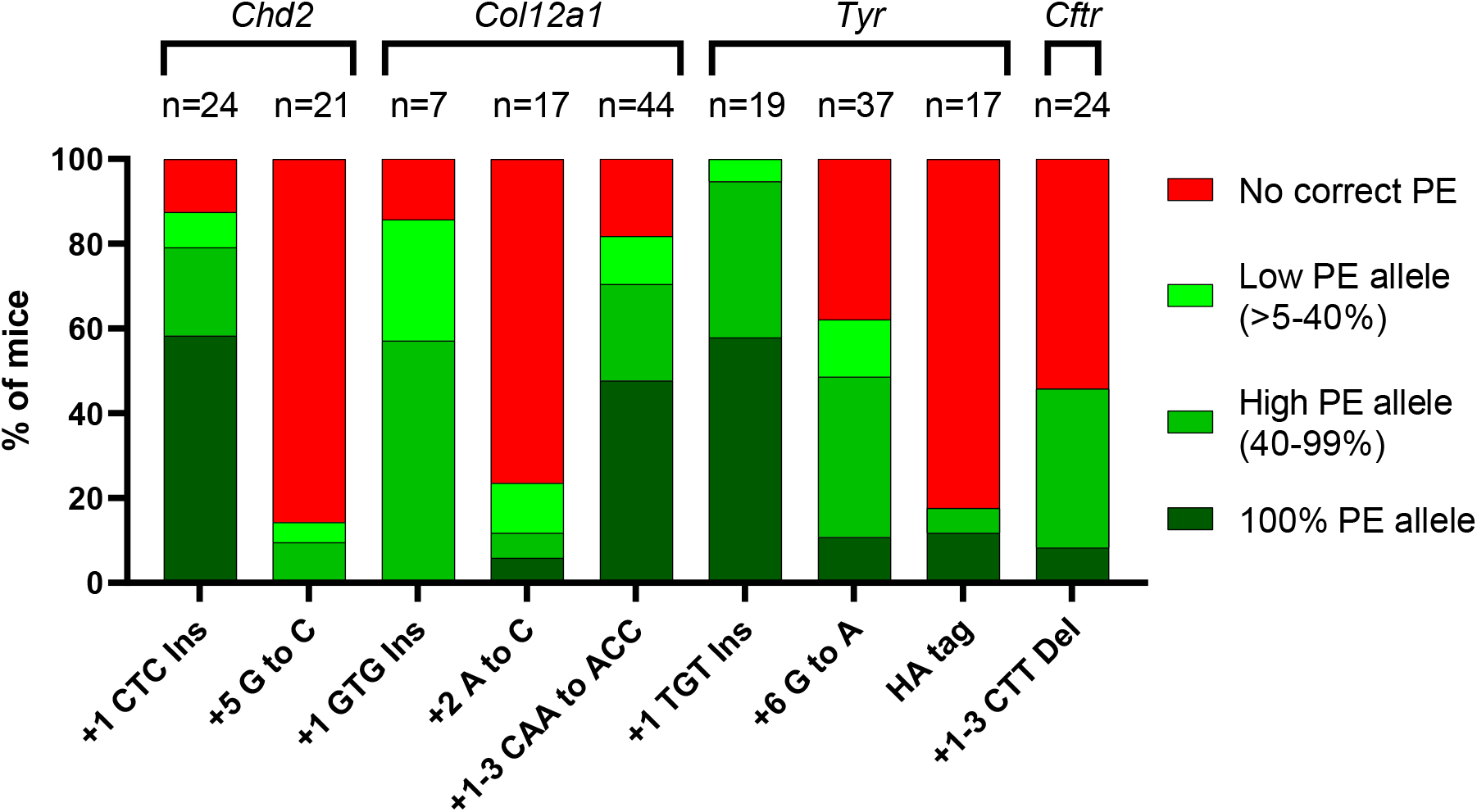
Efficient generation of prime edited mice by zygote injection of nuclease prime editor.

Generation of a +5 G to C single substitution that eliminated the PAM in *Chd2* was relatively inefficient, with only 3 of 21 mice containing the correct edit (Fig. 5). Generating the +2 A to C single substitution in *Col12a1* was also inefficient (Fig. 5) due to re-cutting, as previously observed in mouse ES cells (Supplementary Fig. 6). Remarkably, amending the substitution to avoid re-cutting (+1−3 CAA to ACC) resulted in higher editing efficiency, with 36 of 44 mice (81.8%) carrying the correct edits (Fig. 5). Generating the +6 G to A (PAM-eliminating) single substitution in *Tyr* was also highly efficient, with 23 of 37 (62.16%) mice carrying the correct edit (Fig. 5). We also attempted to generate a longer insertion – specifically a 27 nt HA-tag sequence upstream the *Tyr* stop codon. Although successful, this was relatively inefficient, with 3 of 17 mice carrying correct edits (Fig. 5). Finally, we used nuclease prime editor to model the cystic fibrosis deltaF508 allele (*Cftr* +1−3 CTT del) with 11 of 24 mice (45.8%) containing the correct intended mutation (Fig. 5). Notably, in all samples, the remaining alleles (non-correct edits) were indels or partial template duplications, while the unmodified alleles were rarely detected (Supplementary Fig. 7). Together, these data demonstrate that PE-nuclease can readily generate mice with specific genomic edits.

## DISCUSSION

Although it is a relatively new technique, prime editing has been used in many different cell lines and organisms including plants, zebrafish and *Drosophila* [8–20]. In this study we have created versatile plasmids to perform prime editing PE2, PE3 and PE-nuclease system in mammalian cells. These plasmids provide an “all-in-one” system and include a selection marker (puromycin or GFP) for transfected cells. Importantly, PE constructs of interest can be readily designed using PETAL tool and generated using a single-step cloning protocol. Given the simplicity of the system and that the plasmids are freely available for academic researchers, we anticipate this will expand the use of PE for a range of research applications.

The all-in-one system and the selection step ensure the presence of all PE components inside cells to maximize PE events. Indeed, using this optimized approach we found that the PE efficiency is very high in HEK293T cells (mean of 67%), which is higher than previously observed using multiple vectors without selection [1]. This higher efficiency could also be attributed to differences in the promoter that drives the prime editor, the nuclear localization signal, and the transfection method. The PEA1 constructs use the CbH promoter to drive Cas9, unlike the Anzalone et al. construct which used the CMV promoter. PEA1 also uses the nucleoplasmin nuclear localization signal (NLS) at the C terminus of the prime editor as opposed to SV40 NLS. Lastly, we used nucleofection for HEK293T cell transfection as opposed to lipofection. Surprisingly, despite optimized delivery, PE efficiency is modest in K562 cells (mean of 29%) and relatively poor in HeLa cells (mean of 7.2%). This cell-dependent efficiency of PE is presumably affected by the differences in the relative activity of DNA repair pathways between cell lines. Identifying the key factors for PE repair is critical for further improvement of PE efficiency in cell lines, primary cells and *in vivo* for potential therapeutic applications.

We also generated a Cas9-nuclease prime editor to investigate whether a DNA double-strand break could improve PE efficiency, particularly in K562 and HeLa cells. Interestingly, priming and RT extension of the PBS from the single pegRNA was very efficient using nuclease approach. However, instead of incorporation of the desired genetic alteration, partial template duplications (PTDs) predominate, which appear to result from end-joining repair of the reverse-transcribed RT template at the DNA double-strand break (DSB) site (Supplementary Fig. 5). PTDs also occur with the PE3 nickase system, albeit very infrequently. Development of approaches to reduce PTDs and improve perfect resolution of PE such as by increasing resection and blocking end-joining might pave the way for more efficient PE in hard to edit cells such as K562 and HeLa cells using this nuclease prime editor. Apart from the correct PE allele, the nickase prime editor mostly produces unmodified alleles with few indels. In contrast, the nuclease prime editor produces fewer unmodified alleles and higher indels. This feature of the nuclease prime editor may be an advantage for applications where both PE and indels give benefits such as gene inactivation using PE. However, it is predicted that nuclease prime editor will be more prone to creating off-target cleavages compared to the nickase version as Cas9 nickase has been shown to create minimal off-targets [21]. This Cas9 nuclease off-target effect could be reduced by careful design of the gRNAs or by using high-fidelity Cas9 nuclease [3, 4, 22–24].

We have also shown that zygotic injection of nuclease prime editor mRNA and pegRNA can be used to efficiently generate mutant mice with specific edits. Notably, correctly edited mice were generated for all nine modifications from a single injection session. Previous attempts to use the PE2 or PE3 nickase prime editing systems for mutant mouse generation were hindered by low editing efficiency and the generation of unwanted large intervening deletions (for PE3) resulting from dual-nicking [5–7]. Nuclease prime editor successfully circumvents these limitations. The large intervening deletions do not occur since only one gRNA is used. Efficient DNA breaks facilitated by the Cas9 nuclease also induce high frequency of correct PE. When creating 3 bp insertions at the cut sites, we showed that 87.5%, 85.7% and 100% of mice contained the intended edits when targeting *Chd2, Col12a1* and *Tyr* loci, respectively. Generating substitutions was less efficient and hampered by recutting of the correct edited alleles, particularly when the substitutions have close match to the target sequences [3]. Therefore, this phenomenon should be taken into consideration when editing using nuclease prime editor. We also showed that generating larger insertions such as an HA-Tag could be achieved using the nuclease prime editor, albeit with lower efficiency. Although Sanger analyses indicated 100% pure intended alleles amongst those mice, homozygosity remained unknown as large deletions that delete PCR primer binding sites could occur, which lead to alleles with large deletions being excluded from the analyses [25]. Given these encouraging correct editing efficiencies, using nuclease prime editor for generating mutant mice with specific edits is a promising approach and could be a less expensive alternative compared to the conventional approach using ssDNA long oligo donor [26]. In summary, the nuclease prime editor is a valuable addition to the genome editing toolbox that enables precision editing with higher efficiency than the nickase prime editor at some loci.

## METHODS

### Plasmid generation

To generate PEA1-Puro, PDG459 [2] which was derived from plasmid PX459 V2 [27], was modified to contain the H840A mutation from pCMV-PE2 [1] (a gift from David Liu, Addgene plasmid # 132775) by EcoRV and PmlI fragment sub-cloning. The first hU6-gRNA cassette was then replaced with a hU6-pegRNA cassette generated as a gBlock (IDT) by PciI and Acc65I cloning. Another gBlock containing the reverse transcriptase coding region was generated and cloned to the intermediate plasmid as an XcmI and FseI fragment. Some silent nucleotide changes were introduced to remove BbsI restriction sites. The PEA1-nuclease-puro construct was generated by replacing the H840A sequence region of PEA1-Puro with WT Cas9 nuclease sequence of PDG459 using BmgBI fragment sub-cloning. pCMV_T7-PE2-Nuclease was generated by replacing the H840A region of pCMV-PE2 with the nuclease fragment isolated from PEA1-Nuc by sub-cloning using SacI and EcoRI. PEA1-GFP and PEA1-Nuc-GFP were generated by replacing the puromycin regions of PEA1-Puro and PEA1-Nuc-Puro, respectively, with the GFP sequence from PDG458 [2] using PciI and FseI sub-cloning. Plasmid purification was performed using The QIAprep Spin Miniprep Kit (Qiagen). Plasmids generated from this study were deposited to Addgene (# 171991-171997).

### Generation of targeting constructs using one-step golden gate digestion-ligation cloning

For each PEA1 or PEA1-Nuc targeting construct, 3 oligonucleotide pairs were designed for the pegRNA protospacer, the RT template and the second-nick gRNA protospacer with designated overhangs (Supplementary Table 1). For the PEA1-Nuc targeting constructs, the second-nick gRNA protospacer included a sham gRNA sequence that does not target the mouse or human genome. Oligonucleotide pairs were phosphorylated and annealed by mixing 100 pmol of each pair and 0.5 μL T4 PNK (NEB) then incubated at 37°C for 30 minutes, 95°C for 5 minutes and slowly ramped to RT. Annealed oligonucleotides were then diluted 1 in 250.

One-step golden gate digestion-ligation protocol was performed by mixing 100 ng PEA1 or PEA1-Nuc empty construct with the 3 pairs of diluted oligonucleotides (1 ul each), with the addition of 100 μmol of DTT, 10 μmol of ATP, 1 μL of BbsI (NEB), 1 μL of T4 ligase (NEB) and NEB-2 buffer in 20 μL of reaction. The mixture was placed in a thermocycler and cycled 6 times at 37°C for 5 minutes and 16°C for 5 minutes before bacterial transformation (see Supplementary Materials). Plasmids were purified using the QIAprep Spin Miniprep Kit (Qiagen) and the integration of the oligonucleotides pairs was assayed using Bbs1 digestion (plasmids with putative correct integrations remained circular following the enzymatic digestion). PEA1 targeting plasmids were sequenced using primer GGTTTCGCCACCTCTGACTTG to verify the pegRNA sequences (guide and RT template), and primer CACTCCCACTGTCCTTTCCTAATA to verify the second-nick gRNA sequences. Details of the protocol are provided in Supplementary Note 1.

### Cell culture and transfection

HEK293T (ATCC CRL-3216), K562 (ATCC CCL-243) and HeLa (ATCC CCL-2) cell lines were re-authenticated by Cell Bank Australia. HEK293T and HeLa cells were cultured in Dulbecco’s modified Eagle’s Medium (DMEM; Gibco), while K562 cells were cultured in Iscove’s Modified Dulbecco’s medium (IMDM; Sigma). Each media was supplemented with 10% (v/v) fetal calf serum (FCS; Corning), 1x GlutaMAX (Gibco) and 1x Antibiotic Antimycotic Solution (Sigma). R1 mouse embryonic stem cells (ES cells) from Andras Nagy’s laboratory were cultured in DMEM supplemented with 15% FCS, 1000 units/mL LIF (ESGRO, Sigma), 3 μM CHIR99021 (Sigma), 1 μM PD0325901 (Sigma), 1x GlutaMAX, 100 μM non-essential amino acids (Gibco) and 100 μM 2-mercaptoethanol (Sigma). All cells were maintained in humidified incubators at 37°C with 5% CO_2_ and tested negative for mycoplasma.

Transfection was performed using the Neon nucleofection 100 μL kit (Invitrogen) as per manufacturer’s protocol, delivering 8 μg of plasmid DNA per transfection. HEK239T cells were nucleofected at 1100 V, 20 ms and 2 pulses. Nucleofection for K562 cells was carried out at 1450 V, 10 ms and 3 pulses. For HeLa cells, the program was 1005 V, 35 ms and 2 pulses, while mES cells used 1400 V, 10 ms and 3 pulses. The number of cells used per transfection was 1.5 × 10^6^ for HEK293T cells and 1 × 10^6^ for the K562, HeLa and mES cells. Puromycin selection was initiated 24 hours after nucleofection. The puromycin concentration used was 2 μg/mL for HEK293T, K562 and mES cells and 1 μg/mL for HeLa cells. Puromycin selection was applied for 3 days for HEK293T cells and 2 days for K562, HeLa and mES cells, with selection media changed daily. All the untransfected control cells were expected to die after the puromycin treatment to ensure only transfected cells were collected for analysis. Surviving cells were subsequently recovered in normal media (puromycin-free) until around 70% confluency before collection for genomic DNA extraction.

### Mouse zygote injection

All experiments involving animal use were approved by the South Australian Health & Medical Research Institute (SAHMRI) Animal Ethics Committee. To produce nuclease prime editor mRNA, pCMV_T7-PE2-Nuclease plasmid was linearized using PmeI and purified using DNA Clean & Concentrator-5 (Zymo Research). Purified plasmid was subjected to in vitro transcription (IVT) using mMESSAGE mMACHINE T7 ULTRA Transcription Kit (Ambion/Invitrogen). pegRNA was generated by IVT of PCR purified amplicon [28]. The PCR was performed using the primers listed in Supplementary Table 2. In brief, the forward primer contained the T7 promoter sequences and the gRNA sequences, while the reverse primer contained sequences from the RT template. The corresponding PEA1 targeting constructs were used as the PCR template and the PCR amplicon was purified by QIAquick PCR Purification Kit (Qiagen). IVT was performed using the HiScribe T7 Quick High Yield RNA Synthesis Kit (NEB). Nuclease prime editor mRNA and pegRNA were purified using RNeasy Mini Kit (Qiagen). Details of the protocols for generating pegRNA and nuclease prime editor mRNA are available in Supplementary Note 2 & 3.

Nuclease prime editor mRNA (150 ng/μl) and pegRNA (75 ng/μl) were injected into the cytoplasm of C57BL/6J zygotes using a Femtojet microinjector. Surviving zygotes were transferred into pseudo pregnant females. Mice were harvested at E12.5-13.5 for tissue collection, genomic DNA extractions, PCR and Sanger sequencing.

### DNA extraction and sequencing

DNA extraction from cells and mouse tissues was performed using the Roche High Pure PCR Template Preparation Kit, according to the manufacturer’s protocol. For deep amplicon sequencing, PCR was performed with Nextera-tagged primers under standard Phusion or Q5 Polymerase (New England Biolabs). PCR primers used in this study can be found in Supplementary Table 3. NGS was performed by Australia Genome Research Facility (AGRF) using MiSeq Nano System, paired-end 500 cycle.

NGS was analyzed using the R-GENOME PE-Analyzer online tool [29]. The percentage of correct PE and unmodified alleles was collected directly from the PE-Analyzer calculation. The unintended edit percentage was defined as the alleles apart from correct PE and unmodified alleles (WT). Percentage of ‘any intended edits’ and ‘PTDs’ were calculated by filtering the sequences containing the prime binding site and the edit sequences (details in Supplementary Note 4). The percentage of ‘indels’ was obtained from subtracting the percentage of ‘PTDs’ from ‘unintended edits’. Sanger sequencing of mouse samples from microinjections was analyzed using both ICE and DECODR tools. Allele frequency from ICE analysis was normalized to 100%. Some samples were excluded from the calculation due to failure analyses by ICE and DECODR. The number of mouse samples excluded from the calculation was 3, 1, 3, 2 and 2 for *Chd2* +5 G to C, *Col12a1* +1-3 CAA to ACC, *Tyr* +1 TGT ins, *Tyr* HA tag and *Cftr* +1-3 CTT del, respectively.

## Acknowledgements

We thank Drs David Lui and Andrew Anzalone for providing their published data. This study was supported by funding from the Australian CSIRO Synthetic Biology Future Science Platform awarded to F.A. We would like to acknowledge funding from the Emerging Leaders Development Award of Faculty of Health & Medical Science, University of Adelaide. SAGE is supported by Phenomics Australia via the Australian Government National Collaborative Research Infrastructure Strategy (NCRIS) program.

