## Supplementary Information for "Optimized nickase- and nuclease-based prime editing in human and mouse cells"

Supplementary Figure 1. Sequence view of hU6-pegRNA and hU6-gRNA (second-nick) cassettes and the golden gate cloning sites in PEA1 construct.

#### Benchling sequence view of hU6-pegRNA cassette of PEA1

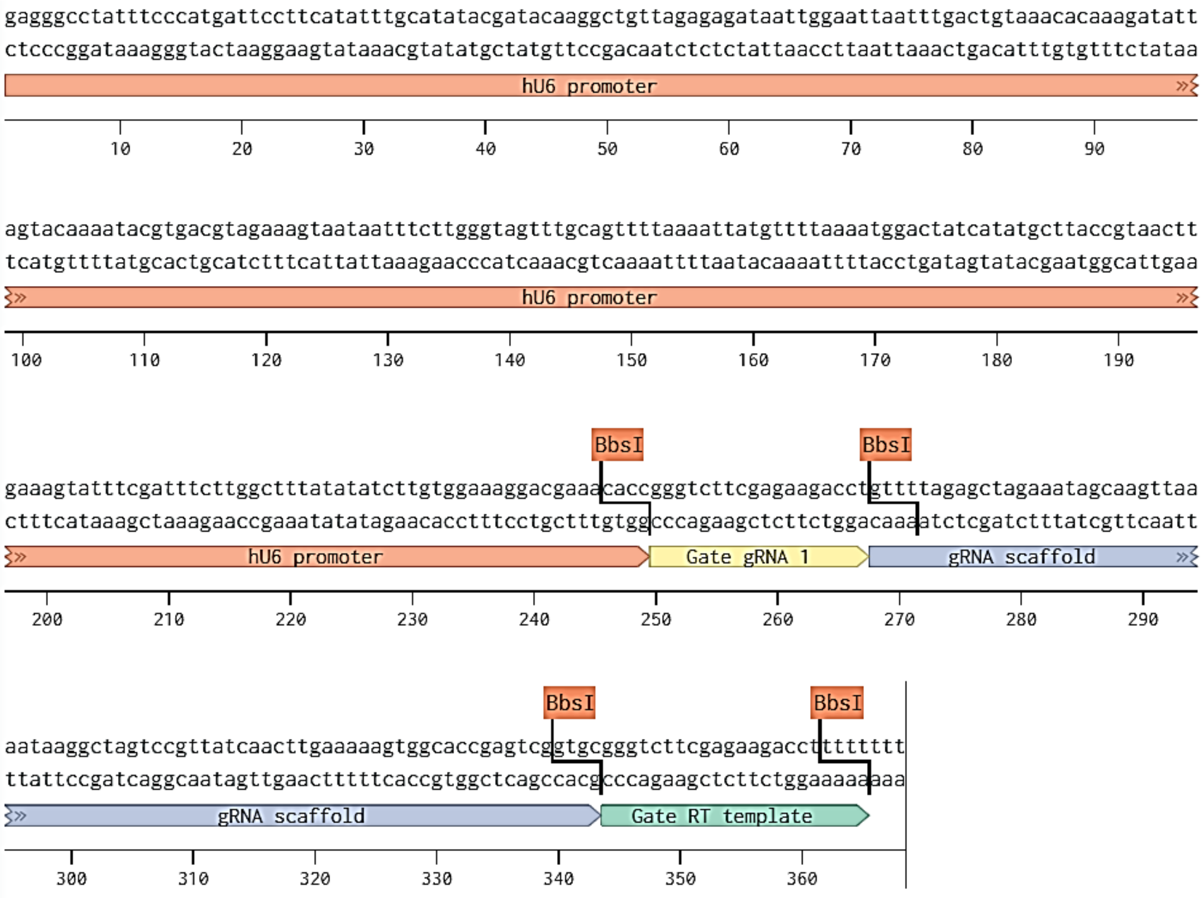

#### Benchling sequence view of hU6-gRNA (second-nick) cassette of PEA1

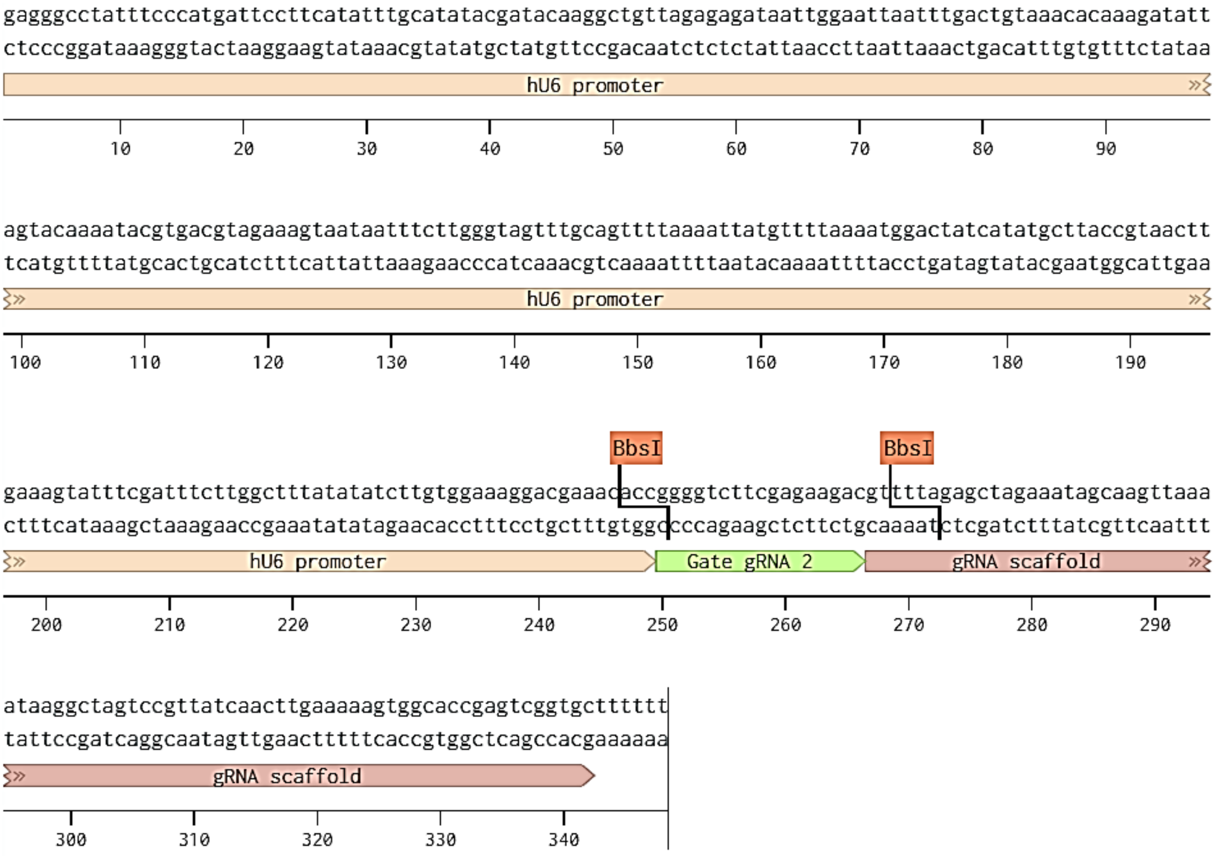

Supplementary Figure 2. Prime Editing Target Locator (PETAL) – an online tool for designing prime editing applications. As input, PETAL takes the target sequence to be edited (top) as well as the final edited sequence (bottom, new inserted sequence highlighted in blue). PETAL identifies all valid guide sequences within the target sequence (top, edited sequence highlighted in dark grey). Users are then able to select the desired pegRNA (blue) and Second Nick (yellow) sequences. PETAL then provides the necessary oligo sequences for both gRNA's as well as those required for the template strand (bottom). The density plot enables users to zoom into the regions of interest.

PETaL - Prime Editing Target Locator

A NEW PRIME EDITING TOOL

Wild-Type Sequence

Wild Type Sequence

CTCCACAGTGCATACGTGGGCTCCAACAGGTCCTCTTCCCTCCCAGTCACTGACTAACCCCGGAACCACACAGCTTCCCGTTCTCAGCTCCACAAACTTGGTGCCAAATTCTTCTCCCCTGGGAAGCATCCCTGGACACTTCCCAAAGGACCCCAGTCACTCCAGCCTGTTGGCTGCCGCTCACTTTGATGTCTGCAGGCCAGATGAGGGCTCCAGATGG
CACATTGTCAGAGGGACACACTGTGGCCCCTGTGCCCAGCCCTGGGCTCTCTGTACATGAAGCAACTCCAGTCCCAAATATGTAGCTGTTTGGGAGGTCAGAAATAGGGGGTCCAGGAGCAAACTCCCCCACCCCCTTTCCAAAGCCCATTCCCTCTTTAGCCAGAGCCGGGGTGTGCAGACGGCAGTCACTAGGG

Edited Sequence

Edited Sequence

CTCCACAGTGCATACGTGGGCTCCAACAGGTCCTCTTCCCTCCCAGTCACTGACTAACCCCGGAACCACACAGCTTCCCGTTCTCAGCTCCACAAACTTGGTGCCAAATTCTTCTCCCCTGGGAAGCATCCCTGGACACTTCCCAAAGGACCCCAGTCACTCCAGCCTGTTGGCTGCCGCTCACTTTGATGTCTGCAGGCCAGATGACAAGGGCTCCAG
ATGGCACATTGTCAGAGGGACACACTGTGGCCCCTGTGCCCAGCCCTGGGCTCTCTGTACATGAAGCAACTCCAGTCCCAAATATGTAGCTGTTTGGGAGGTCAGAAATAGGGGGTCCAGGAGCAAACTCCCCCACCCCCTTTCCAAAGCCCATTCCCTCTTTAGCCAGAGCCGGGGTGTGCAGACGGCAGTCACTAGGG

Submit

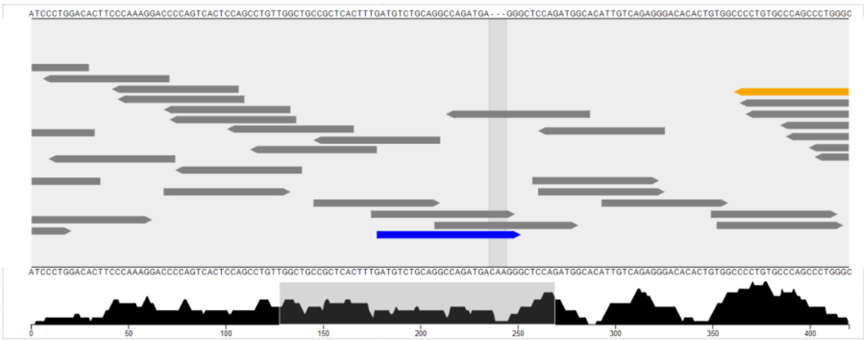

Target Selection and Oligos:

| Guide Type | Name | Sequence |
|---|---|---|
| pegRNA | Guide | GATGTCTGCAGGCCAGATGA |
| pegRNA | Oligo #1 | caccGATGTCTGCAGGCCAGATGA |
| pegRNA | Oligo #2 | aaacTCATCTGGCCTGCAGACATC |
| Second Nick | Guide | gAGAGCCCAGGGCTGGGCACA |
| Second Nick | Oligo #1 | taaaacTGTGCCCAGCCCTGGGCTCT |
| Second Nick | Oligo #2 | accgAGAGCCCAGGGCTGGGCACAgt |
| Template | Guide | CATCTGGAGCCCTTGTCATCTGGCCTGC |
| Template | Oligo #1 | gtgcCATCTGGAGCCCTTGTCATCTGGCCTGC |
| Template | Oligo #2 | aaaaGCAGGCCAGATGACAAGGGCTCCAGATG |

Supplementary Figure 3. Example of Bbs1 check digest of plasmids resulting from one-step digestion-ligation cloning to generate PEA1-Puro *VEGFA* +4 C ins. All plasmids except for plasmid 2 and plasmid 6 had complete integration of the oligo pairs. Plasmid 6 seemed to lack one pair of oligo duplex integration and therefore could be digested and produced linear plasmid.

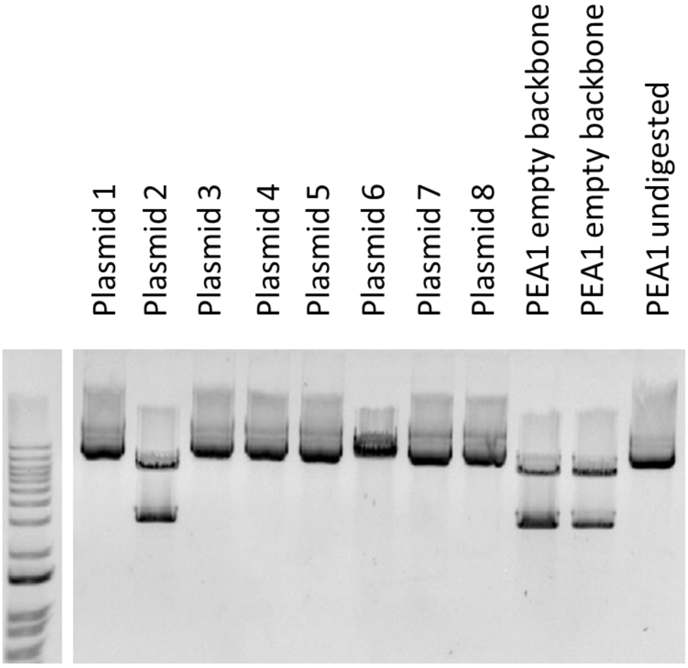

Supplementary Figure 4. Partial template duplications were also observed as unintended editing outcomes of PE3 in HEK293T cells.

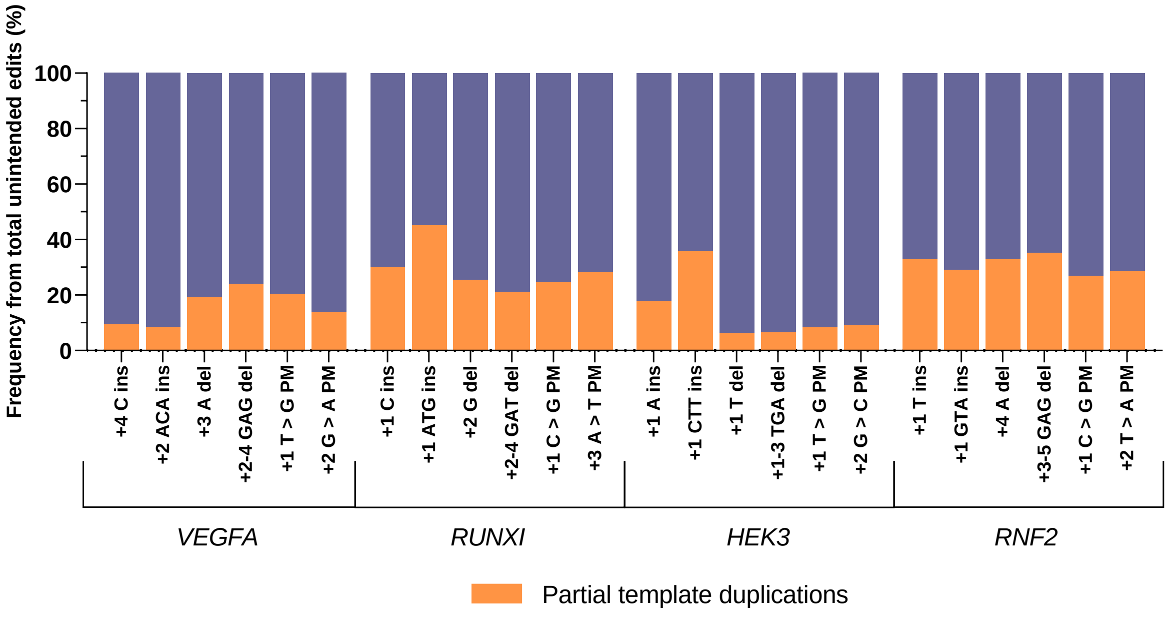

Supplementary Figure 5. Mechanism of partial template duplication (PTD) events.

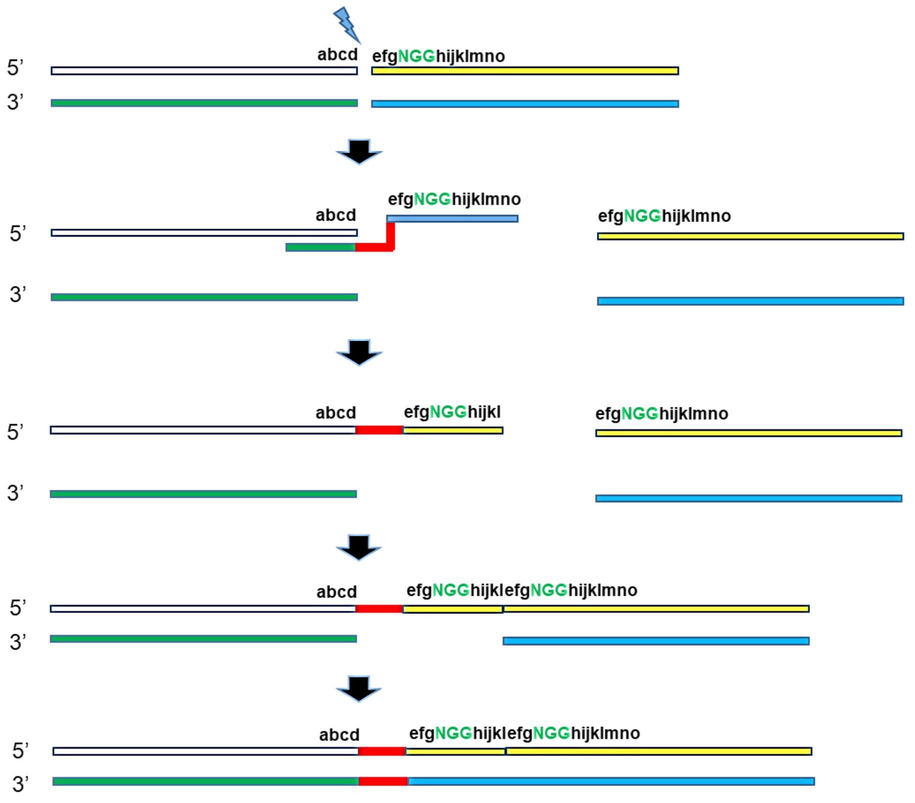

DSB by nuclease prime editor + pegRNA

RT-template of pegRNA (PBS) anneals with the edited strand.

Reverse transcription, genomic DNA copies the RT-template. Edited strand contains ssDNA containing the edits (red) and the homology sequences.

Instead of annealing with the homology counterpart, this ssDNA undergoes end-joining with the DNA break junction.

Extension of the non-edited strand (blue) and ligation to join the DNA, generating PTD end product.

Supplementary Figure 6. Examples of edits found in mouse ES cells targeted with PEA1-Nuc Col12a1 +2 A to C that indicate re-cutting events after prime editing events.

```
TGCCCTGATTAAACCTATTGGAAGAGCTTGACTTCCATGGTTCCACAATGGGTCCATTATGTGCTGGGCTGGGCCCTGTTGTTTTAT
||||||||||||||||||||||||||||||||||           |.||||||||||||||||||||||||||||||||||||||||
TGCCCTGATTAAACCTATTGGAAGAGCTTGACTT-----------CCATGGGTCCATTATGTGCTGGGCTGGGCCCTGTTGTTTTAT
```

14.37%
(n=3)

```
TGCCCTGATTAAACCTATTGGAAGAGCTTGACTTCCATGGTTCCACAATGGGTCCATTATGTGCTGGGCTGGGCCCTGTTGTTTTAT
||||||||||||||||||||||||||||||||||||||||||   |.||||||||||||||||||||||||||||||||||||||||
TGCCCTGATTAAACCTATTGGAAGAGCTTGACTTCCATGGTT---CCATGGGTCCATTATGTGCTGGGCTGGGCCCTGTTGTTTTAT
```

2.77%
(n=3)

Supplementary Figure 7. Frequency of PE, indels, PTD and WT alleles in individual mice generated by nuclease prime editor.

### indicates 3 or more different alleles were detected in this individual mouse.
Alleles with 2 or more bp non-correct insertions were classified as PTDs.

PE-Nuc mice *Chd2* +1 CTC ins

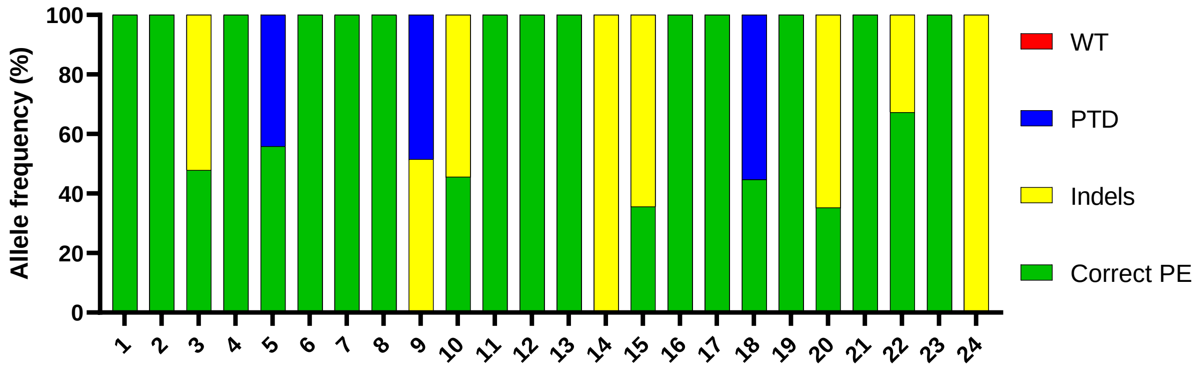

PE-Nuc mice *Chd2* +5 G to C

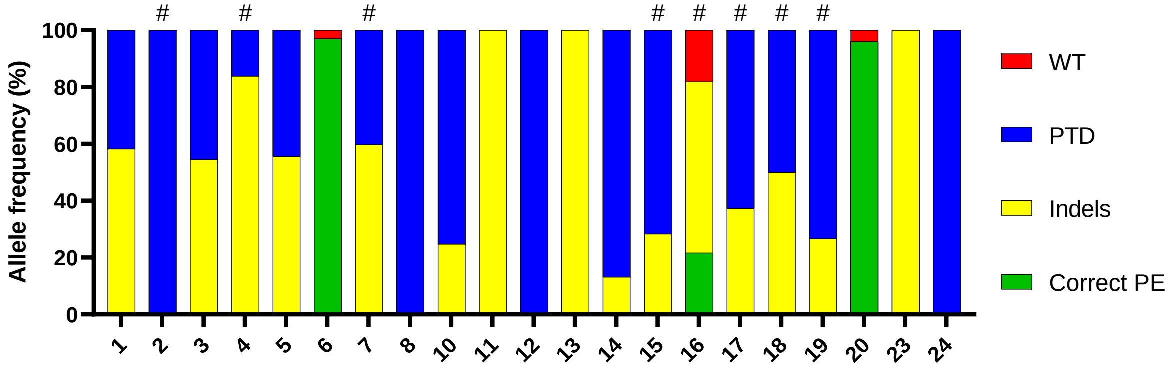

PE-Nuc mice *Col12a1* +1 GTG ins

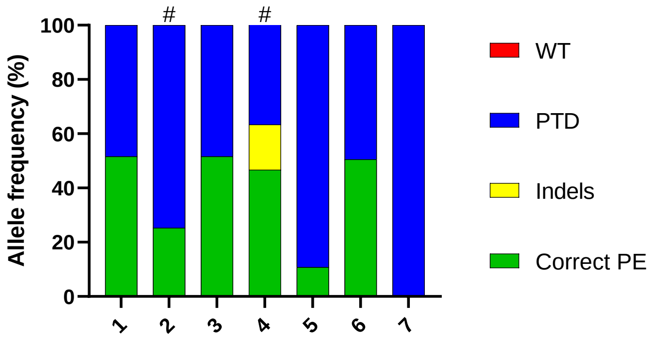

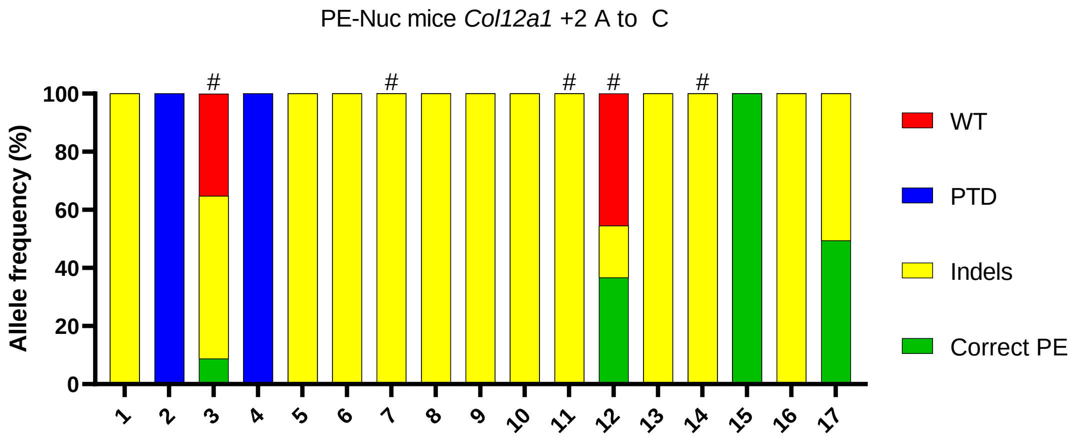
PE-Nuc mice Col12a1 +2 A to C
Allele frequency (%)
WT
PTD
Indels
Correct PE

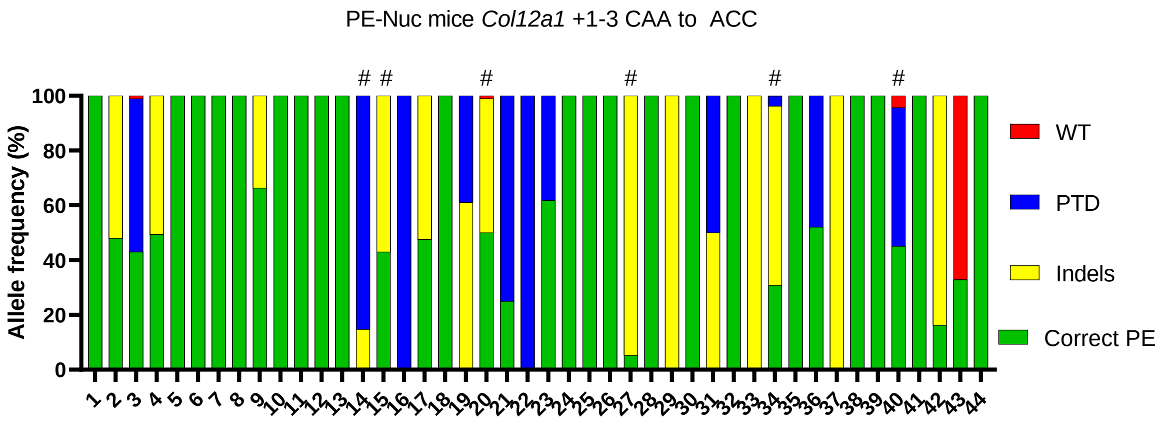
PE-Nuc mice Col12a1 +1-3 CAA to ACC
Allele frequency (%)
WT
PTD
Indels
Correct PE

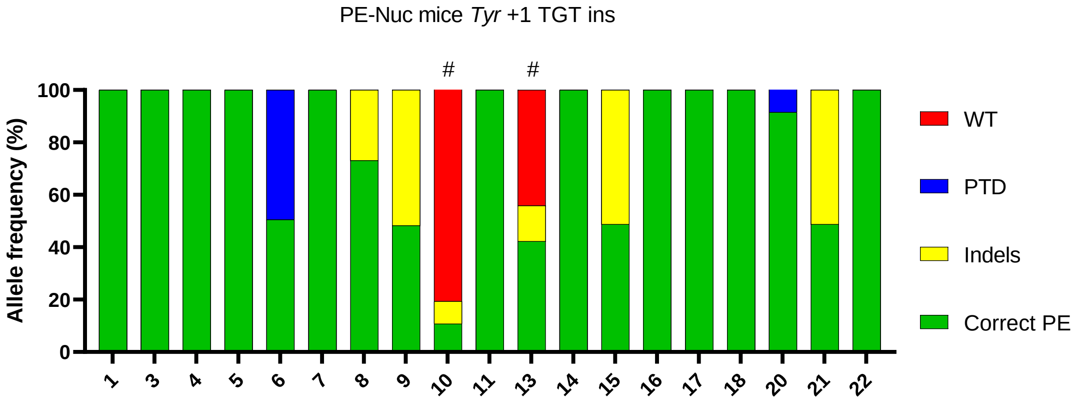
PE-Nuc mice Tyr +1 TGT ins
Allele frequency (%)
WT
PTD
Indels
Correct PE

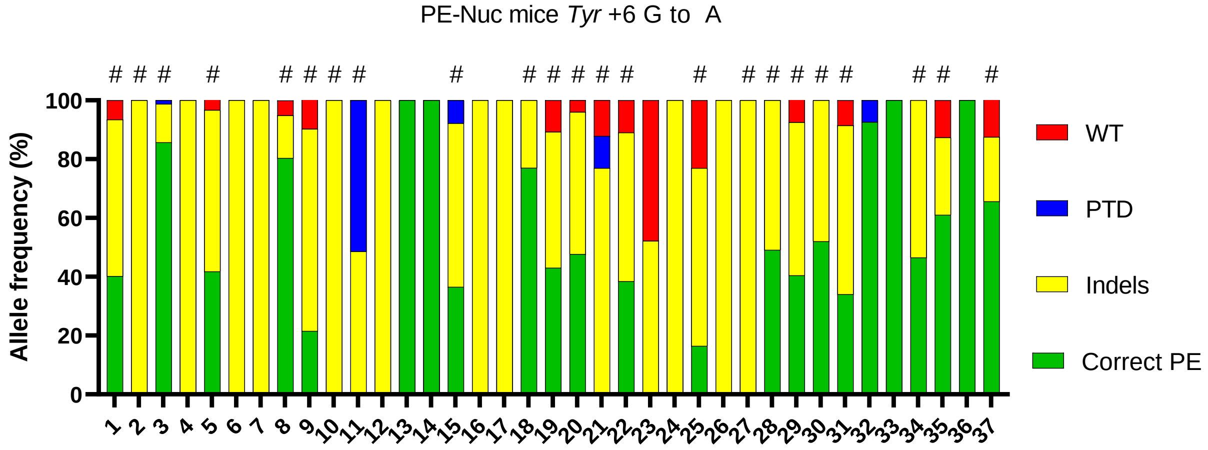
PE-Nuc mice Tyr +6 G to A
Allele frequency (%)
WT
PTD
Indels
Correct PE

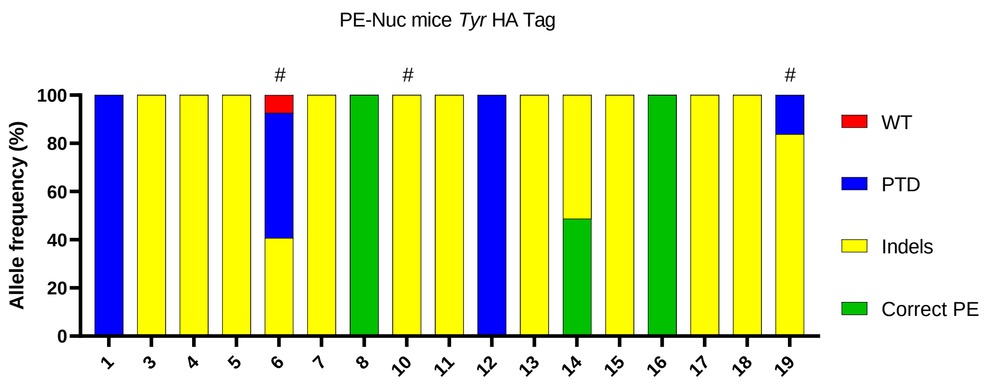

PE-Nuc mice *Tyr* HA Tag
Allele frequency (%)
0
20
40
60
80
100
#
#
#
1
3
4
5
6
7
8
WT
PTD
Indels
Correct PE

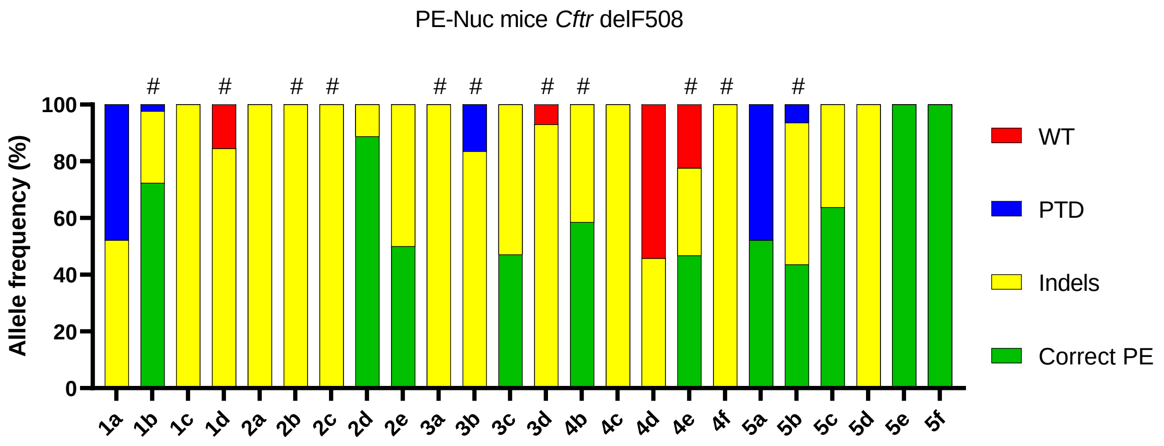

PE-Nuc mice *Cftr* delF508
Allele frequency (%)
0
20
40
60
80
100
#
#
#
#
#
#
#
#
#
#
#
1a
1b
1c
1d
2a
2b
2c
2d
2e
3a
3b
3c
3d
4b
4c
4d
4e
4f
5a
5b
5c
5d
5e
5f
WT
PTD
Indels
Correct PE

Supplementary Table 1. Oligo lists to generate the PEA1 targeting constructs.

| PEA1 targeting construct | Oligo pair 1 (gRNA) | Oligo pair 2 (repair template) | Oligo pair 3 (second-nick gRNA) |
|---|---|---|---|
| *HEK3* +1 A ins | caccGGCCCAGACTGAGCACGTGA | gtgcTCTGCCATCATCGTGCTCAGTCTG | accgTCAACCAGTATCCCGGTGCgt |
| | aaacTCACGTGCTCAGTCTGGGCC | aaaaCAGACTGAGCACGATGATGGCAGA | taaaacGCACCGGGATACTGGTTGA |
| *HEK3* +1 CTT ins | caccGGCCCAGACTGAGCACGTGA | gtgcTCTGCCATCAAAGCGTGCTCAGTCTG | accgTCAACCAGTATCCCGGTGCgt |
| | aaacTCACGTGCTCAGTCTGGGCC | aaaaCAGACTGAGCACGCTTTGATGGCAGA | taaaacGCACCGGGATACTGGTTGA |
| *HEK3* +1 T del | caccGGCCCAGACTGAGCACGTGA | gtgcTCTGCCATCCGTGCTCAGTCTG | accgTCAACCAGTATCCCGGTGCgt |
| | aaacTCACGTGCTCAGTCTGGGCC | aaaaCAGACTGAGCACGGATGGCAGA | taaaacGCACCGGGATACTGGTTGA |
| *HEK3* +1-3 TGA del | caccGGCCCAGACTGAGCACGTGA | gtgcTGGAGGAAGCAGGGCTTCCTTTCCTCTGCCACGTGCTCAGTCTG | accgTCAACCAGTATCCCGGTGCgt |
| | aaacTCACGTGCTCAGTCTGGGCC | aaaaCAGACTGAGCACGTGGCAGAGGAAAGGAAGCCCTGCTTCCTCCA | taaaacGCACCGGGATACTGGTTGA |
| *HEK3* +1 T to G PM | caccGGCCCAGACTGAGCACGTGA | gtgcTCTGCCATCCCGTGCTCAGTCTG | accgTCAACCAGTATCCCGGTGCgt |
| | aaacTCACGTGCTCAGTCTGGGCC | aaaaCAGACTGAGCACGGGATGGCAGA | taaaacGCACCGGGATACTGGTTGA |
| *HEK3* +2 G to C | caccGGCCCAGACTGAGCACGTGA | gtgcTCTGCCATGACGTGCTCAGTCTG | accgTCAACCAGTATCCCGGTGCgt |
| | aaacTCACGTGCTCAGTCTGGGCC | aaaaCAGACTGAGCACGTCATGGCAGA | taaaacGCACCGGGATACTGGTTGA |
| *RNF2* +1 T ins | caccGTCATCTTAGTCATTACCTG | gtgcAACGAACACCTCAGAGTAATGACTAAGATG | accgTCAACCATTAAGCAAAACATgt |
| | aaacCAGGTAATGACTAAGATGAC | aaaaCATCTTAGTCATTACTCTGAGGTGTTCGTT | taaaacATGTTTTGCTTAATGGTTGA |
| *RNF2* +1 GTA ins | caccGTCATCTTAGTCATTACCTG | gtgcAACGAACACCTCAGTACGTAATGACTAAGATG | accgTCAACCATTAAGCAAAACATgt |
| | aaacCAGGTAATGACTAAGATGAC | aaaaCATCTTAGTCATTACGTACTGAGGTGTTCGTT | taaaacATGTTTTGCTTAATGGTTGA |
| *RNF2* +4 A del | caccGTCATCTTAGTCATTACCTG | gtgcAACGAACACCCAGGTAATGACTAAGATG | accgTCAACCATTAAGCAAAACATgt |
| | aaacCAGGTAATGACTAAGATGAC | aaaaCATCTTAGTCATTACCTGGGTGTTCGTT | taaaacATGTTTTGCTTAATGGTTGA |
| *RNF2* +3-5 GAG del | caccGTCATCTTAGTCATTACCTG | gtgcAACGAACACAGGTAATGACTAAGATG | accgTCAACCATTAAGCAAAACATgt |
| | aaacCAGGTAATGACTAAGATGAC | aaaaCATCTTAGTCATTACCTGTGTTCGTT | taaaacATGTTTTGCTTAATGGTTGA |
| *RNF2* +1 C to G | caccGTCATCTTAGTCATTACCTG | gtgcAACGAACACCTCACGTAATGACTAAGATG | accgTCAACCATTAAGCAAAACATgt |
| | aaacCAGGTAATGACTAAGATGAC | aaaaCATCTTAGTCATTACGTGAGGTGTTCGTT | taaaacATGTTTTGCTTAATGGTTGA |
| *RNF2* +2 T to A | caccGTCATCTTAGTCATTACCTG | gtgcAACGAACACCTCTGGTAATGACTAAGATG | accgTCAACCATTAAGCAAAACATgt |
| | aaacCAGGTAATGACTAAGATGAC | aaaaCATCTTAGTCATTACCAGAGGTGTTCGTT | taaaacATGTTTTGCTTAATGGTTGA |
| *RUNX1* +1 C ins | caccGCATTTTCAGGAGGAAGCGA | gtgcTGTCTGAAGCCATCGGCTTCCTCCTGAAAAT | accgATGAAGCACTGTGGGTACGAgt |
| | aaacTCGCTTCCTCCTGAAAATGC | aaaaATTTTCAGGAGGAAGCCGATGGCTTCAGACA | taaaacTCGTACCCACAGTGCTTCAT |
| *RUNX1* +1 ATG ins | caccGCATTTTCAGGAGGAAGCGA | gtgcTGTCTGAAGCCATCCATGCTTCCTCCTGAAAAT | accgATGAAGCACTGTGGGTACGAgt |
| | aaacTCGCTTCCTCCTGAAAATGC | aaaaATTTTCAGGAGGAAGCATGGATGGCTTCAGACA | taaaacTCGTACCCACAGTGCTTCAT |
| *RUNX1* +2 G del | caccGCATTTTCAGGAGGAAGCGA | gtgcTGTCTGAAGCCATGCTTCCTCCTGAAAAT | accgATGAAGCACTGTGGGTACGAgt |
| | aaacTCGCTTCCTCCTGAAAATGC | aaaaATTTTCAGGAGGAAGCATGGCTTCAGACA | taaaacTCGTACCCACAGTGCTTCAT |
| *RUNX1* +2-4 GAT del | caccGCATTTTCAGGAGGAAGCGA | gtgcTGTCTGAAGCCGCTTCCTCCTGAAAAT | accgATGAAGCACTGTGGGTACGAgt |
| | aaacTCGCTTCCTCCTGAAAATGC | aaaaATTTTCAGGAGGAAGCGGCTTCAGACA | taaaacTCGTACCCACAGTGCTTCAT |
| *RUNX1* +1 C to G | caccGCATTTTCAGGAGGAAGCGA | gtgcTGTCTGAAGCCATCCCTTCCTCCTGAAAAT | accgATGAAGCACTGTGGGTACGAgt |
| | aaacTCGCTTCCTCCTGAAAATGC | aaaaATTTTCAGGAGGAAGGGATGGCTTCAGACA | taaaacTCGTACCCACAGTGCTTCAT |

| | | | |
|---|---|---|---|
| *RUNX1* +3 A to T | caccGCATTTTCAGGAGGAAGCGA | gtgcTGTCTGAAGCCAACGCTTCCTCCTGAAAAT | accgATGAAGCACTGTGGGTACGAgt |
| | aaacTCGCTTCCTCCTGAAAATGC | aaaaATTTTCAGGAGGAAGCGTTGGCTTCAGACA | taaaacTCGTACCCACAGTGCTTCAT |
| *VEGFA* +4 C ins | caccGATGTCTGCAGGCCAGATGA | gtgcAATGTGCCATCTGGAGCCGCTCATCTGGCCTGCAGA | accgATGTACAGAGAGCCCAGGGCgt |
| | aaacTCATCTGGCCTGCAGACATC | aaaaTCTGCAGGCCAGATGAGCGGCTCCAGATGGCACATT | taaaacGCCCTGGGCTCTCTGTACAT |
| *VEGFA* +2 ACA ins | caccGATGTCTGCAGGCCAGATGA | gtgcAATGTGCCATCTGGAGCCCTTGTCATCTGGCCTGCAGA | accgATGTACAGAGAGCCCAGGGCgt |
| | aaacTCATCTGGCCTGCAGACATC | aaaaTCTGCAGGCCAGATGACAAGGGCTCCAGATGGCACATT | taaaacGCCCTGGGCTCTCTGTACAT |
| *VEGFA* +3 A del | caccGATGTCTGCAGGCCAGATGA | gtgcAATGTGCCATCTGGAGCCCCATCTGGCCTGCAGA | accgATGTACAGAGAGCCCAGGGCgt |
| | aaacTCATCTGGCCTGCAGACATC | aaaaTCTGCAGGCCAGATGGGGCTCCAGATGGCACATT | taaaacGCCCTGGGCTCTCTGTACAT |
| *VEGFA* +2-4 GAG del | caccGATGTCTGCAGGCCAGATGA | gtgcAATGTGCCATCTGGAGCCATCTGGCCTGCAGA | accgATGTACAGAGAGCCCAGGGCgt |
| | aaacTCATCTGGCCTGCAGACATC | aaaaTCTGCAGGCCAGATGGCTCCAGATGGCACATT | taaaacGCCCTGGGCTCTCTGTACAT |
| *VEGFA* +1 T to G | caccGATGTCTGCAGGCCAGATGA | gtgcAATGTGCCATCTGGAGCCCTCCTCTGGCCTGCAGA | accgATGTACAGAGAGCCCAGGGCgt |
| | aaacTCATCTGGCCTGCAGACATC | aaaaTCTGCAGGCCAGAGGAGGGCTCCAGATGGCACATT | taaaacGCCCTGGGCTCTCTGTACAT |
| *VEGFA* +2 G to A | caccGATGTCTGCAGGCCAGATGA | gtgcAATGTGCCATCTGGAGCCCTTATCTGGCCTGCAGA | accgATGTACAGAGAGCCCAGGGCgt |
| | aaacTCATCTGGCCTGCAGACATC | aaaaTCTGCAGGCCAGATAAGGGCTCCAGATGGCACATT | taaaacGCCCTGGGCTCTCTGTACAT |
| *Chd2* +1 CTC ins | caccGCGGTAGCTCCCAGAACGGT | gtgcGATGCCCACCgagGTTCTGGGAGCTA | accgACCATCAGTATGAGCAGCATgt |
| | aaacACCGTTCTGGGAGCTACCGC | aaaaTAGCTCCCAGAACctcGGTGGGCATC | taaaacATGCTGCTCATACTGATGGT |
| *Chd2* +5 G to C | caccGCGGTAGCTCCCAGAACGGT | gtgcGATGCgCACCGTTCTGGGAGCTA | accgACCATCAGTATGAGCAGCATgt |
| | aaacACCGTTCTGGGAGCTACCGC | aaaaTAGCTCCCAGAACGGTGcGCATC | taaaacATGCTGCTCATACTGATGGT |
| *Col12a1* +1 GTG ins | caccGACTTCCATGGTTCCACAA | gtgcAATGGACCCATTGcacTGGAACCATGGAA | accgCCTGAGCAGGCCACGAACAgt |
| | aaacTTGTGGAACCATGGAAGTC | aaaaTTCCATGGTTCCAgtgCAATGGGTCCATT | taaaacTGTTCGTGGCCTGCTCAGG |
| *Col12a1* +2 A to C | caccGACTTCCATGGTTCCACAA | gtgcAATGGACCCATgGTGGAACCATGGAA | accgCCTGAGCAGGCCACGAACAgt |
| | aaacTTGTGGAACCATGGAAGTC | aaaaTTCCATGGTTCCACcATGGGTCCATT | taaaacTGTTCGTGGCCTGCTCAGG |
| *Col12a1* +1-3 CAA to ACC | caccGACTTCCATGGTTCCACAA | gtgcAATGGACCCAggtTGGAACCATGGAA | accgGGCAGCGCGGCTATCGTGGCgt |
| | aaacTTGTGGAACCATGGAAGTC | aaaaTTCCATGGTTCCAaccTGGGTCCATT | taaaacGCCACGATAGCCGCGCTGCC |
| *Tyr* +1 TGT ins | caccGCAAAAGAATGCTGCCCACCA | gtgcATCACCCATCCATGGacaTGGGCAGCATTCT | accgCACTGGACAGAAGGATATCCgt |
| | aaacTGGTGGGCAGCATTCTTTTGC | aaaaAGAATGCTGCCCAtgtCCATGGATGGGTGAT | taaaacGGATATCCTTCTGTCCAGTG |
| *Tyr* +6 G to A | caccGCAAAAGAATGCTGCCCACCA | gtgcATCACCCATtCATGGTGGGCAGCATTCT | accgCACTGGACAGAAGGATATCCgt |
| | aaacTGGTGGGCAGCATTCTTTTGC | aaaaAGAATGCTGCCCACCATGaATGGGTGAT | taaaacGGATATCCTTCTGTCCAGTG |
| *Tyr* HA-Tag | caccGTTTCCTAGGATGTTCACAGA | gtgcTCAGAGCCATCTgTACCCATACGATGTTCCAGATTACGCTtaaGTGAACATCCTAG | accgGGCAGCGCGGCTATCGTGGCgt |
| | aaacTCTGTGAACATCCTAGGAAAC | aaaaCTAGGATGTTCACttaAGCGTAATCTGGAACATCGTATGGGTAcAGATGGCTCTGA | taaaacGCCACGATAGCCGCGCTGCC |
| *Mixl1* +1 CTT ins (Nick +48) | caccGCAAGTGGATGTCTGGGTACA | gtgcTCCGACAGACCATGTaagACCCAGACATCCAC | accgCAAGCGCACGTCGTTCAGCTgt |
| | aaacTGTACCCAGACATCCACTTGC | aaaaGTGGATGTCTGGGTcttACATGGTCTGTCGGA | taaaacAGCTGAACGACGTGCGCTTG |
| *Mixl1* +1 A to G (Nick +48) | caccGCAAGTGGATGTCTGGGTACA | gtgcTCCGACAGACCATGcACCCAGACATCCAC | accgCAAGCGCACGTCGTTCAGCTgt |
| | aaacTGTACCCAGACATCCACTTGC | aaaaGTGGATGTCTGGGTgCATGGTCTGTCGGA | taaaacAGCTGAACGACGTGCGCTTG |
| *Mixl1* +1-3 ACA del (Nick +48) | caccGCAAGTGGATGTCTGGGTACA | gtgcTCCGACAGACCAACCCAGACATCCAC | accgCAAGCGCACGTCGTTCAGCTgt |
| | aaacTGTACCCAGACATCCACTTGC | aaaaGTGGATGTCTGGGTTGGTCTGTCGGA | taaaacAGCTGAACGACGTGCGCTTG |

| | | | |
|---|---|---|---|
| *Mixl1* +1 CTT ins (Nick -60) | caccGCAAGTGGATGTCTGGGTACA | gtgcTCCGACAGACCATGTaagACCCAGACATCCAC | accgCTACCCGAGTCCAGGATCCgt |
| | aaacTGTACCCAGACATCCACTTGC | aaaaGTGGATGTCTGGGTcttACATGGTCTGTCGGA | taaaacGGATCCTGGACTCGGGTAG |
| *Mixl1* +1 A to G (Nick -60) | caccGCAAGTGGATGTCTGGGTACA | gtgcTCCGACAGACCATGcACCCAGACATCCAC | accgCTACCCGAGTCCAGGATCCgt |
| | aaacTGTACCCAGACATCCACTTGC | aaaaGTGGATGTCTGGGTgCATGGTCTGTCGGA | taaaacGGATCCTGGACTCGGGTAG |
| *Mixl1* +1-3 ACA del (Nick -60) | caccGCAAGTGGATGTCTGGGTACA | gtgcTCCGACAGACCAACCCAGACATCCAC | accgCTACCCGAGTCCAGGATCCgt |
| | aaacTGTACCCAGACATCCACTTGC | aaaaGTGGATGTCTGGGTTGGTCTGTCGGA | taaaacGGATCCTGGACTCGGGTAG |
| *EphB2* loxP site 1 (R) | caccGCCATGGTCTCAGGTAATAGC | gtgcTTGTCTCAGCTCCTGCTATAACTTCGTATAATGTATGCTATACGAAGTTATcaattgATTACCTGAGACCA | accgAGAGAAAGATGAGACTGGAgt |
| | aaacGCTATTACCTGAGACCATGGC | aaaaTGGTCTCAGGTAATcaattgATAACTTCGTATAGCATACATTATACGAAGTTATAGCAGGAGCTGAGACAA | taaaacTCCAGTCTCATCTTTCTCT |
| *EphB2* loxP site 2 (L2) | caccGCAGTCACTCTGTAAACCCTG | gtgcGAAGAGCGCGACCCCAGATAACTTCGTATAATGTATGCTATACGAAGTTATgatatcGGTTTACAGAGTGA | accgAGTATGGAGCAGAGAGGCTgt |
| | aaacCAGGGTTTACAGAGTGACTGC | aaaaTCACTCTGTAAACCgatatcATAACTTCGTATAGCATACATTATACGAAGTTATCTGGGGTCGCGCTCTTC | taaaacAGCCTCTCTGCTCCATACT |
| *EphB2* loxP site 3 (L3) | caccGCCAAGAGCCTAGGCAATCGT | gtgcAGAGGTAGACTCCCACGATAACTTCGTATAATGTATGCTATACGAAGTTATgatatcATTGCCTAGGCTCT | accgCCACTCCACCAGTAAAGAAAgt |
| | aaacACGATTGCCTAGGCTCTTGGC | aaaaAGAGCCTAGGCAATgatatcATAACTTCGTATAGCATACATTATACGAAGTTATCGTGGGAGTCTACCTCT | taaaacTTTCTTTACTGGTGGAGTGG |
| *Cftr* +1-3 CTT del | caccATCAAAGAAAATATCATCTT | gtgcATCATAGGAAACACCAATGATATTTTCTTTG | accgGGCAGCGCGGCTATCGTGGCgt |
| | aaacAAGATGATATTTTCTTTGAT | aaaaCAAAGAAAATATCATTGGTGTTTCCTATGAT | taaaacGCCACGATAGCCGCGCTGCC |

gRNA sequences are underlined. Red highlight indicates extra G was added to the gRNA sequences. Oligo pair 3 for PEA1-Nuc targeting constructs used a sham targeting oligos which are the same oligos highlighted in blue).

Supplementary Table 2. Primers to generate IVT template of pegRNAs for mouse zygote injections.

| Target | Forward primer | Reverse primer |
|---|---|---|
| Chd2 +1 CTC ins | TTAATACGACTCACTATAGGCGGTAGCTCCCAGAACGGT | aaaaTAGCTCCCAGAACctcGGTGGGCATC |
| Chd2 +5 G to C | TTAATACGACTCACTATAGGCGGTAGCTCCCAGAACGGT | aaaaTAGCTCCCAGAACGGTGcGCATC |
| Col12a1 +1 GTG ins | TTAATACGACTCACTATAGTGACTTCCATGGTTCCACAA | aaaaTTCCATGGTTCCAgtgCAATGGGTCCATT |
| Col12a1 +2 A to C | TTAATACGACTCACTATAGTGACTTCCATGGTTCCACAA | aaaaTTCCATGGTTCCACcATGGGTCCATT |
| Col12a1 +1-3 CAA to ACC | TTAATACGACTCACTATAGTGACTTCCATGGTTCCACAA | aaaaTTCCATGGTTCCAaccTGGGTCCATT |
| Tyr +1 TGT ins | TTAATACGACTCACTATAGCAAAAGAATGCTGCCCACCA | aaaaAGAATGCTGCCCAtgtCCATGGATGGGTGAT |
| Tyr +6 G to A | TTAATACGACTCACTATAGCAAAAGAATGCTGCCCACCA | aaaaAGAATGCTGCCCACCATGaATGGGTGAT |
| Tyr HA-Tag | TTAATACGACTCACTATAGTTTCCTAGGATGTTCACAGA | aaaaCTAGGATGTTCACttaAGCGTAATCTGGAACATCGTATGGGTAcAGATGGCTCTGA |
| Cftr +1-3 CTT del | TTAATACGACTCACTATAGATCAAAGAAAATATCATCTT | aaaaCAAAGAAAATATCATTGGTGTTTCCTATGAT |

T7 promoter sequences are highlighted in green. The reverse primers are the same as the bottom primers used for oligo pair 2 for generating PEA1 targeting constructs.

Supplementary Table 3. List of PCR primers for sequencing.

| Target sites | Forward | Reverse |
|---|---|---|
| *HEK3* | GGGAAACGCCCATGCAATTA | CAGAGATCAACCAGATTACCCCA |
| *RNF2* | ACGTAGGAATTTTGGTGGGACA | ACAGATGTAGCACCAACCATGGA |
| *RUNX1* | AGAGAGATGTAGGGCTAGAGGG | CACTTGACAAAGTTCTCACGC |
| *VEGFA* | CTCCACAGTGCATACGTGGG | CCCTAGTGACTGCCGTCTG |
| *Chd2* | CTTGCAGATCGAGGAGACTGG | CTCTCCTGCATCCTCAGGCT |
| *Col12a1* | CAGTATGAAGTCATGTGCGGTC | CAATGGAAGACAGGAGTAGGGC |
| *Tyr* | GTCTGTGACACTCATTAACCTATTGGTGC | TCAACTGCGGAAACTGTAAGTTTGGA |
| *Tyr*-HA Tag | GGAGCTGTTATTGCTGCAGCTC | ACCAGCTCAATTAGTTGTAAGAGG |
| *Mixl1* | CCGCTTTCCCCATCTTCC | GACTTCCCAGCACCTCCACT |
| *EphB2* LoxP site 1 | AGGTAGGCACCACCATGATC | AGGCTGGCATGGGTTAGTTC |
| *EphB2* LoxP site 1 | GACCACTCCACCAGTAAAGAAAGG | CAAGCAGGATATGAGGGAGCAG |
| *EphB2* LoxP site 1 | GGCAGGTGGATCTCTGAGTTTG | CCACCCTGTGCTATCTATCAGTCA |
| *Cftr* | TCACAGCAATTTAAGTAGGGGC | GGGATGATACCGTCCATCTTGG |

For NGS PCRs, primers contain Nextera adapter sequences at the 5’ end. The adapter sequences for the forward primer are TCGTCGGCAGCGTCAGATGTGTATAAGAGACAG. The adapter sequences for the reverse primer are GTCTCGTGGGCTCGGAGATGTGTATAAGAGACAG.

**Supplementary Note 1. One-step digestion-ligation protocol using PEA1 to generate PE targeting constructs.**

Oligos for guide and RT template insertion into plasmid need to be of the following form:

pegRNA guide:

```
5'-CACCGNNNNNNNNNNNNNNNNNNNN-3'
    3'-CNNNNNNNNNNNNNNNNNNNNCAAA-5'
```

pegRNA RT template

```
5'-GTGCNNNNNNNNNNNNNNNNNNNNNNNNNNNNNN-3'
    3'-NNNNNNNNNNNNNNNNNNNNNNNNNNNNNNAAAA-5'
```

Second-nick guide

```
5'-ACCGNNNNNNNNNNNNNNNNNNNNGT-3'
    3'-NNNNNNNNNNNNNNNNNNNNCAAAAT-5'
```

If the first N on the top strand for each guide is a G, it should be excluded.

- For the guides, the N's (typically 20 bases) in the two top strands comprise the guide sequence, which target the identical gRNA binding sequences followed by PAMs in the genomic DNA.
- The overhangs allow the oligos to bind the complementary overhanging DNA at the cut sites in the plasmid created by *Bbs*I digestion.
- The U6 promoter is more efficient if it starts transcription with a G, this is the reason for the extra G/C in the first pair of oligos, this doubles as the first base in the guide which is why it should be excluded if the guide starts with a G. The G is also present as part of the overhang in the second pair of oligos.
- The extra GT/CA in the second pair of oligos completes the gRNA scaffold.

1. Mix the following reagents in a PCR tube for each of the two inserts:

| Reagent | | Amount |
|---|---|---|
| MQ $H_2O$ | | 6.5 µL |
| NEB T4 DNA Ligase Buffer with 10 mM ATP | (10x) | 1 µL |
| top oligo | (100 µM) | 1 µL |
| bottom oligo | (100 µM) | 1 µL |
| NEB T4 PNK | (10 U/µL) | 0.5 µL |
| **Total** | | **10 µL** |

2. Place each mixtures in thermocycler with the following parameters:

| | | |
|---|---|---|
| **1** | 37 °C | 30 min |
| **2** | 95 °C | 5 min |
| **3** | Ramp to 25 °C @ 0.1 °C/s | ∞ |

3. Dilute the 3 sets of phospho-annealed oligos 1:250 with MQ $H_2O$ in a 1.5 mL tube.

| Reagent | Amount |
|---|---|
| MQ $H_2O$ | 249 µL |
| phospho-annealed oligo | 1 µL |
| **Total** | **250 µL** |

4. Mix the following reagents in a PCR tube:

| Reagent | | Amount |
|---|---|---|
| MQ $H_2O$ | | 10 µL |
| PEA1 empty plasmid | (100 ng/µL) | 1 µL |
| phospho-annealed oligo pair 1 (1:250) | | 1 µL |
| phospho-annealed oligo pair 2 (1:250) | | 1 µL |
| phospho-annealed oligo pair 3 (1:250) | | 1 µL |
| NEBuffer 2.1 | (10x) | 2 µL |
| DTT | (10 mM) | 1 µL |
| ATP | (10 mM) | 1 µL |
| NEB *Bbs*I | (5 U/µL) | 1 µL |
| T4 DNA Ligase | (400 U/µL) | 1 µL |
| **Total** | | **20 µL** |

Note: We showed that removing NEBuffer 2.1, DTT, ATP components and replacing it with 2 ul of 10x T4 Ligase Buffer (add extra MQ to reach total volume of 20 ul) could lead to successful reaction.

5. Place in thermocycler with the following parameters:

| | | |
|---|---|---|
| **1** | 37 °C | 5 min |
| **2** | 16 °C | 5 min |
| **3** | Go to step 1 | 5 times |

6. Transform to competent cells.
   Recommended: Incubating the reaction overnight at 4 C before transformation could lead to higher number of colonies.

7. Check digest using Bbs1. Plasmids with complete integrations remain circular.
8. Sequence verify using primers GGTTTCGCCACCTCTGACTTG and CACTCCCACTGTCCTTTCCTAATA.

**Supplementary Note 2. Generation of pegRNA protocol.**

Kit: NEB HiScribe™ T7 Quick High Yield RNA Synthesis.

> T7 guide primer for PCR of gRNA oligo needs to be of the following form:
> 5’-TTAATACGACTCACTATAGNNNNNNNNNNNNNNNNNNN-3’
> Where the N’s are identical to your specific guide.
> Reverse primer for PCR of gRNA oligo needs to be:
> 5’- AAAANNNNNNNNNNNNNNNNNNNNNNNNNN-3’
> Where the N’s are the RT template sequence.
> This oligos are the same oligos used for RT template bottom oligos in the digestion-ligation protocol.

1. Generate the relevant PEA1 targeting plasmid (miniprep)
2. Mix the following master mix reagents in a 1.5 mL tube and aliquot into 6 PCR tubes:

| Reagent | Amount | 6x MM |
|---|---|---|
| MQ $H_2O$ | 12.3 µL | 73.8 µL |
| NEB Phusion HF Reaction Buffer (5x) | 4 µL | 24 µL |
| T7 guide primer (10 µM) | 1 µL | 6 µL |
| Reverse primer (10 µM) | 1 µL | 6 µL |
| NTP Mix (10 mM) | 0.5 µL | 3 µL |
| NEB Phusion HF DNA Polymerase (2 U/µL) | 0.2 µL | 1.2 µL |
| 1 µL PEA1 targeting plasmid (~1-3 ng/µL) | 1 | 6 |
| **Total** | **20 µL** | **120 µL** |

3. Place the tubes in a thermocycler with the following parameters:

| | | |
|---|---|---|
| **1** | 98 °C | 3 min |
| **2** | 98 °C | 15 s |
| **3** | 60 °C | 20 s |
| **4** | 72 °C | 15 s |
| **5** | Go to step 2 | 32 times |
| **6** | 72 °C | 5 min |
| **7** | 4 °C | ∞ |

4. Make a 1% agarose gel and run 5 µL of the PCR products

> Testing the plasmid has the correct insert.

> Band should be present at ~100 bp.

5. Combine all PCR reactions and perform **Qiagen PCR Purification** in a single column.
6. Use NanoDrop to measure concentration of DNA.

> Confirms the DNA is still present.

7. Perfrom IVT by mixing the following reagents in a PCR tube:

| Reagent | Amount |
|---|---|
| Nuclease-free MQ $H_2O$ | up to 40 µL |
| NEB NTP Buffer Mix (20 mM) | 20 µL |
| Purified PCR product | ~1000 ng |
| NEB T7 RNA Polymerase Mix | 4 µL |
| **Total** | **~40 µL** |

Note: half reaction (total 20 ul) is also possible.

8. Incubate O/N @ 37 °C in thermocycler.

Note: 3 hours is also possible.

9. Transfer 2 µL to PCR tube for testing later.
10. Mix the following reagents in a PCR tube:

| Reagent | Amount |
|---|---|
| Nuclease-free MQ $H_2O$ | 60 µL |
| IVT gRNA product | 40 µL |
| NEB DNase I (RNase-free) (2 U/µL) | 4 µL |
| **Total** | **104 µL** |

Degrades DNA.

11. Incubate 15 min @ 37 °C.
12. Transfer 2 µL to PCR tube for testing later.
13. Perform **Qiagen RNEasy Mini Kit RNA Cleanup**, eluting in 30 µL.
14. Check RNA on gel (RNAse free technique should be applied).

**Supplementary Note 3. Generation of nuclease prime editor mRNA protocol.**

- Linearize plasmid PE2-Nuc using Pme1
  - MQ = X ul
  - Cut smart buffer = 6 ul
  - Plasmid = Y ul (10 ug)
  - Pme1 = 3 ul
  - Total 60 ul
  - Incubate 37 C for 2 hours
- Purify the linearized plasmid using Zymo DNA clean and concentrator 5
  - Add 200 ul binding buffer
  - Spin
  - Add 200 ul wash buffer spin
  - Repeat wash
  - Add 12 ul RNAse-free water
  - Spin
- Setup IVT using Mmessage ultra kit
  - T7 Arca = 10 ul
  - Buffer = 2 ul
  - Linearized plasmid = X ul (1.5-2 ug)
  - T7 enzyme = 2 ul
  - RNAse-free water = Y ul (total 20 ul)
  - Incubate 37 C for 3 hours
  - Add 1 ul of DNAse, incubate 30 min 37 C
  - Add 36 ul water + 20 ul EPAP + 10 ul MnCl2 + 10 ul ATP (all included in the kit)
  - Take 2.5 ul for gel checking
  - Add 4 ul of EPAP enzyme, incubate 37 C for 20 min
  - Keep the reaction on ice
  - Take 2.5 ul for gel checking
  - Proceed to RNA clean up using RNAeasy kit (elute in 35 ul of water).
- Zygote microinjection mix
  - MQ = X ul
  - 10x injection buffer = 1.5 ul
  - Nuclease prime editor mRNA = Y ul (final 150 ng/ul)
  - pegRNA = Z ul (final 75 ng/ul)
  - Total = 15 ul

10X injection buffer

| Reagent | | Amount |
|---|---|---|
| pH 8.0 EDTA | (0.5 M) | 10 µL |
| pH 7.5 Tris | (1 M) | 500 µL |
| Nuclease-free MQ $H_2O$ | | 4.49 mL |
| **Total** | | **5 mL** |

(Filtered into aliquots in 1.5 mL tubes and stored @ - 20 °C.)

**Supplementary Note 4. Data analysis using Rgenome PE-Analyzer.**

**%Correct PE**
Correct PE can be directly gathered from the generated analysis

| Prime editing |
| --- |
| 64 (1.4%) |

**%Unmodified (WT)**
WT frequency can be gathered directly by clicking the WT column. Ensure you don't filter any sequence here.

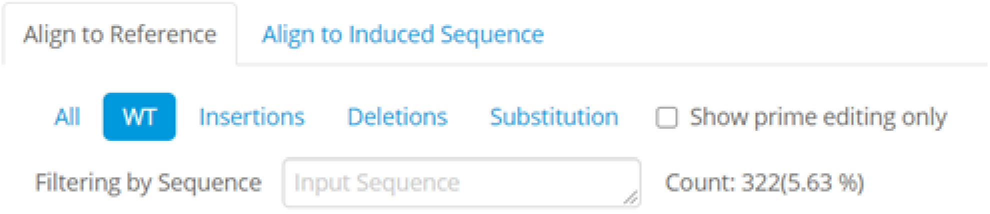

**%Unintended edits = 100 - %Correct PE - %WT**

**%Any intended edits:** any alleles containing prime edited sequences (modified and/or unmodified)
To get total PE, filter sequences starting from PBS to 2nt of the edit.

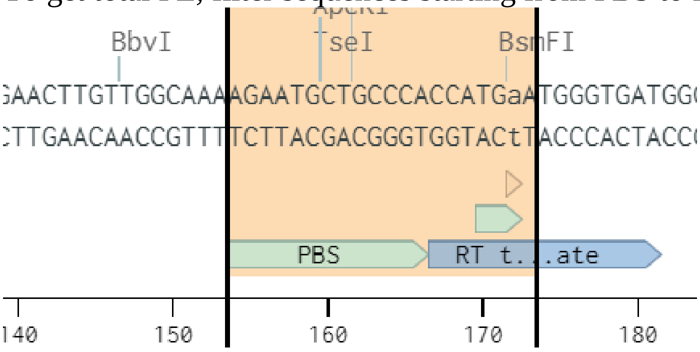

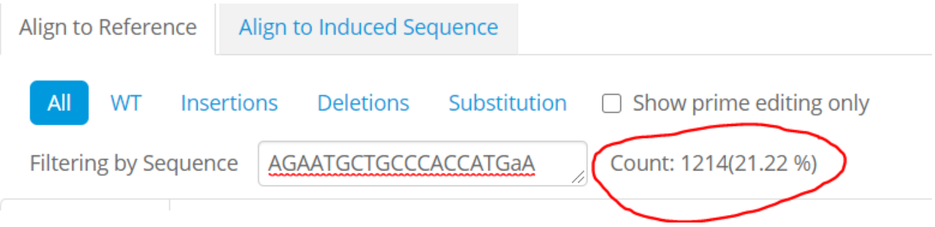

The sequence list to filter to get total PE:
HEK3:
+1 A ins: CAGACTGAGCACGaT
+1 CTT ins: CAGACTGAGCACGct
+1 T del: CAGACTGAGCACGGA
+1-3 TGA del: CAGACTGAGCACGTGG
+1 T to G PM: CAGACTGAGCACGgG
+2 G to C PM: CAGACTGAGCACGTcA

**RNF2:**
+1 T ins: GaGTAATGACTAAGATG
+1 GTA ins: acGTAATGACTAAGATG
+4 A del: CCCAGGTAATGACTAAGATG
+3-5 GAG del: ACAGGTAATGACTAAGATG
+1 C to G PM: AcGTAATGACTAAGATG

+2 T to A PM: CtGGTAATGACTAAGATG

**RUNX1:**
+1 C ins: ATTTTCAGGAGGAAGcC
+1 ATG ins: ATTTTCAGGAGGAAGCat
+2 G del: ATTTTCAGGAGGAAGCAT
+2-4 GAT del: ATTTTCAGGAGGAAGCGG
+1 C to G PM : ATTTTCAGGAGGAAGgG
+3 A to T PM : ATTTTCAGGAGGAAGCGtT

**VEGFA:**
+4 C ins:TCTGCAGGCCAGATGAGcG
+2 ACA ins: TCTGCAGGCCAGATGac
+3 A del: TCTGCAGGCCAGATGGG
+2-4 GAG del: TCTGCAGGCCAGATGG
+1 T to G PM: TCTGCAGGCCAGAgG
+2 G to A PM: TCTGCAGGCCAGATaA

Mouse ES cells
Mixl1 +1 CTT ins: agACCCAGACATCCAC
Mixl1 +1-3 ACA del : CAACCCAGACATCCAC
Mixl1 +1 A to G : GcACCCAGACATCCAC

Tyr +1 TGT ins : AGAATGCTGCCCAtg
Tyr +6 G to A : AGAATGCTGCCCACCATGaA

Chd2 +1 CTC ins : agGTTCTGGGAGCTA
Chd2 +5 G to C : CgCACCGTTCTGGGAGCTA

Col12a1 +1 GTG ins : TTCCATGGTTCCAgt
Col12a1 +2 A to C : TTCCATGGTTCCACcA

**%Any loxP:** any alleles containing loxP sequences (modified and/or unmodified)
LoxP site 1 ATAACTTCGTATAATGTATGCTATACGAAGTTATcaattg
LoxP site 2 ATAACTTCGTATAATGTATGCTATACGAAGTTATgatatc
LoxP site 3 ATAACTTCGTATAATGTATGCTATACGAAGTTATgatatc

**%Partial template duplications (PTDs)**
To get the frequency of PTDs, filter the same sequences above into the "insertions" column.

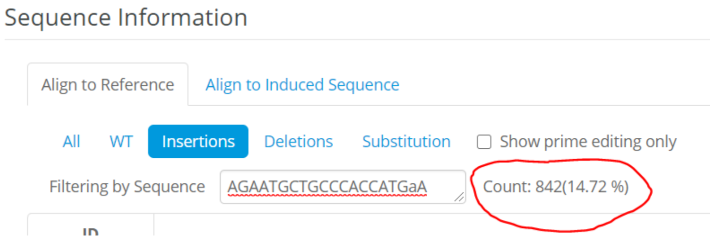

If the edit is substitution or deletion, collect the frequency of PTD straight away from the count.

If the edit is insertion, the PTD = the count – the correct PE.

**%Indels = %unintended edits - %PTDs**
